# Mechanisms of spontaneous Ca^2+^ release-mediated arrhythmia in a novel 3D human atrial myocyte model: I. Transverse-axial tubule variation

**DOI:** 10.1101/2022.07.15.500273

**Authors:** X. Zhang, H. Ni, S. Morotti, C.E.R. Smith, D. Sato, W.E. Louch, A.G. Edwards, E. Grandi

## Abstract

Intracellular calcium (Ca^2+^) cycling is tightly regulated in the healthy heart ensuring effective contraction. This is achieved by transverse (t)-tubule membrane invaginations that facilitate close coupling of key Ca^2+^-handling proteins such as the L-type Ca^2+^ channel (LCC) and Na^+^-Ca^2+^ exchanger (NCX) on the cell surface with ryanodine receptors (RyRs) on the intracellular Ca^2+^ store. Though less abundant and regular than in the ventricle, t-tubules also exist in atrial myocytes as a network of transverse invaginations with axial extensions known as the transverse-axial tubule system (TATS). In heart failure and atrial fibrillation there is TATS remodeling that is associated with aberrant Ca^2+^-handling and Ca^2+^-induced arrhythmic activity, however the mechanism underlying this is not fully understood. To address this, we developed a novel 3D human atrial myocyte model that couples electrophysiology and Ca^2+^-handling with variable TATS organization and density. We extensively parameterized and validated our model against experimental data to build a robust tool examining TATS regulation of subcellular Ca^2+^ release. We found that varying TATS density and thus the localization of key Ca^2+^-handling proteins has profound effects on Ca^2+^ handling. Following TATS loss there is reduced NCX that results in increased cleft Ca^2+^ concentration through decreased Ca^2+^ extrusion. This elevated Ca^2+^ increases RyR open probability causing spontaneous Ca^2+^ releases and promotion of arrhythmogenic waves (especially in the cell interior) that leads to voltage instabilities through delayed afterdepolarizations. In summary, this study demonstrates a mechanistic link between TATS remodeling and Ca^2+^-driven proarrhythmic behavior that likely reflects the arrhythmogenic state observed in disease.

**Key Points:**

- Transverse-axial tubule systems (TATS) modulate Ca^2+^ handling and excitation-contraction coupling in atrial myocytes, with TATS remodeling in heart failure and atrial fibrillation associated with altered Ca^2+^ cycling and subsequent arrhythmogenesis.
- To investigate the poorly understood mechanisms linking TATS variation and spontaneous Ca^2+^ release, we built, parameterized and validated a 3D human atrial myocyte model coupling electrophysiology and spatially-detailed subcellular Ca^2+^ handling governed by the TATS.
- Simulated TATS loss causes diastolic Ca^2+^ and voltage instabilities through reduced NCX-mediated Ca^2+^ removal, cleft Ca^2+^ accumulation and increased RyR open probability, resulting in spontaneous Ca^2+^ release and promotion of arrhythmogenic waves and delayed afterdepolarizations.
- At fast electrical rates typical of atrial tachycardia/fibrillation, spontaneous Ca^2+^ releases are larger and more frequent in the cell interior than at the periphery.
- Our work provides mechanistic insight into how atrial TATS remodeling can lead to Ca^2+^- driven instabilities that may ultimately contribute to the arrhythmogenic state in disease.

## Introduction

Intracellular Ca^2+^ is an important determinant of both contractile and electrophysiologic function of cardiac myocytes through the process of excitation-contraction coupling (ECC). Dysregulation of Ca^2+^ cycling is known to be the primary driver of arrhythmia in specific diseases or drug-responses (e.g., catecholaminergic polymorphic ventricular tachycardia, digoxin toxicity), and also contributes to more common arrhythmogenic conditions with broader pathophysiologic changes, such as atrial fibrillation (AF) and heart failure (HF) (Denham *et al.*, 2018). It has long been known that disease-induced ionic remodeling can lead to abnormal Ca^2+^ handling (Denham *et al.*, 2018); however, subcellular structural remodeling can also contribute to the observed alterations. Transverse tubules (TTs) are invaginations of the sarcolemma that play a key role in ECC in ventricular myocytes by closely (~10 nm) juxtaposing sarcolemmal voltage-gated L-type Ca^2+^ channels (LCCs) with ryanodine receptors (RyRs) on the membrane of the sarcoplasmic reticulum (SR) to facilitate rapid and synchronous Ca^2+^-induced Ca^2+^ release (CICR) (Cheng *et al.*, 1993; Bers, 2002), and efficient fluid exchange between TT lumens and extracellular areas (Hong *et al.*, 2014; Rog-Zielinska *et al.*, 2021). TTs were historically thought to be absent in atrial myocytes (Hüser *et al.*, 1996; Greiser *et al.*, 2014) thus limiting triggered CICR to the cell periphery with release in the cell interior reliant on propagation through recruitment of the inner RyRs (Blatter *et al.*, 2003). More recently, however, several studies have demonstrated an extensive TT network in the atria of a variety of species, though less dense and organized, and more variable than in ventricles. Interestingly, in contrast to the ventricle, evidence suggests that longitudinally-oriented axial tubules may be prominent in atrial myocytes and provide extensive transverse-axial tubule (TAT) extensions of sarcolemmal invaginations that also couple RyRs with membrane ion channels (Arora *et al.*, 2017; Brandenburg *et al.*, 2018, 2019). Indeed, several reports have revealed the existence of a transverse-axial tubular system (TATS) in atrial myocytes, with varying density and organization in different cardiac regions (left/right chamber, endo-/epicardium) (Kirk *et al.*, 2003; Frisk *et al.*, 2014; Glukhov *et al.*, 2015; Arora *et al.*, 2017) and species, e.g., mouse (Yue *et al.*, 2017; Brandenburg *et al.*, 2018), rat (Kirk *et al.*, 2003; Rasmussen *et al.*, 2004; Woo *et al.*, 2005; Dibb *et al.*, 2009; Trafford *et al.*, 2013; Frisk *et al.*, 2014; Glukhov *et al.*, 2015; Brandenburg *et al.*, 2018), canine (Dolber *et al.*, 1994; Melnyk *et al.*, 2002; Trafford *et al.*, 2013), rabbit (Greiser *et al.*, 2014; Brandenburg *et al.*, 2018), pig (Frisk *et al.*, 2014; Gadeberg *et al.*, 2016; Brandenburg *et al.*, 2018), cow (Richards *et al.*, 2011), horse (Richards *et al.*, 2011), sheep (Dibb *et al.*, 2009; Lenaerts *et al.*, 2009; Caldwell *et al.*, 2014), and human (Richards *et al.*, 2011; Brandenburg *et al.*, 2018).

The importance of the TATS in atrial cells is highlighted by the impact of its remodeling in disease. TATS alterations are known to be a major contributor to AF (Trafford *et al.*, 2013), combining with ionic remodeling to destabilize the bidirectional interaction between electrical activation and Ca^2+^ signaling in atrial myocytes. Currently, limitations in existing experimental methods make it difficult to separate the independent contributions of ionic and structural remodeling at the cellular level. Furthermore, while clear, though variable, TATS are seen in human tissue (Richards *et al.*, 2011), only sparse TATS have been reported in isolated human atrial myocytes (Greiser *et al.*, 2014), most likely due to membrane damage during enzymatic digestion (Chen *et al.*, 2015). These challenges limit our mechanistic and quantitative understanding of the precise role of TATS variability in health and TATS remodeling in human AF. Indeed, while disruption of TATS architecture is associated with arrhythmia in numerous states (e.g., in ventricular myocytes in HF, and myocardial infarction (Louch *et al.*, 2006)), a direct mechanistic link between remodeling of atrial myocyte ultrastructure and arrhythmogenesis has not yet been established. In this context, mathematical models are powerful tools to both fill the gaps in the available data sources and reveal the independent contributions of TATS remodeling. Over the last decade, computational models have been developed including detailed spatial and temporal characteristics of myocyte Ca^2+^ signaling, accounting for thousands of stochastic Ca^2+^ release units, connected with the global (whole-cell) electrophysiologic behavior (Shiferaw *et al.*, 2005; Song *et al.*, 2015). These sophisticated models have been employed to study how varying cell structural properties affect local and global atrial Ca^2+^ signaling, ECC, and the development of alternans and spontaneous Ca^2+^ release (SCRs) (Colman *et al.*, 2016; Shiferaw *et al.*, 2017, 2018, 2020; Marchena & Echebarria, 2018, 2020; Sutanto *et al.*, 2018). Experimentally, spontaneous Ca^2+^ release (SCR) and delayed afterdepolarizations (DADs) have been shown to precede the development of classical markers of ionic remodeling in AF (Voigt *et al.*, 2014), and while RyR hyperactivity is clearly involved (Voigt *et al.*, 2012, 2014), computational analyses suggest that complete detubulation could provide an additional mechanism for increased SCR and DADs during Ca^2+^ overload (Li *et al.*, 2012). Though simulations of perturbed TATS structure have been performed, they have typically been heuristic in their approach, with limited coupling to experiments and sporadic robust validation of the coupling of voltage and Ca^2+^ dynamics.

To quantitatively understand the mechanistic link between variation in human atrial ultrastructure and myocyte function and arrhythmogenesis, we built an integrative modeling framework coupling our common-pool model of the human atrial myocyte action potential (AP) (**Fig. 1Ai**) (Grandi *et al.*, 2011; Morotti *et al.*, 2016b) with a three-dimensional (3D) model of subcellular Ca^2+^ signaling (**Fig. 1Aii**) based on a rabbit ventricular myocyte (Restrepo *et al.*, 2008; Sato & Bers, 2011), with varying randomly generated TATS (**Fig. 1AiIi**) utilizing human atrial myocyte data (Brandenburg *et al.*, 2018). We first present our integrative model parameterization and validation against broad independent functional datasets, illustrating local and global Ca^2+^ handling and AP properties in the atria, and then interrogate the model to explain the relationship between the TATS, SCRs, and DADs in human atrial myocytes. We show that reducing TATS density leads to local increases in cleft Ca^2+^ concentration through reduced Ca^2+^ removal by the Na^+^-Ca^2+^ exchanger (NCX). This Ca^2+^ elevation drives cleft Ca^2+^-dependent increases in RyR open probability (P_O_) and subsequent SCRs that demonstrate how atrial TATS remodeling can lead to Ca^2+^-driven proarrhythmic behavior. In a companion paper (Zhang et al), we used this robustly validated framework to study the independent effects of varying key Ca^2+^ handling protein expression and distribution on SCRs and DADs.

**Figure 1.**
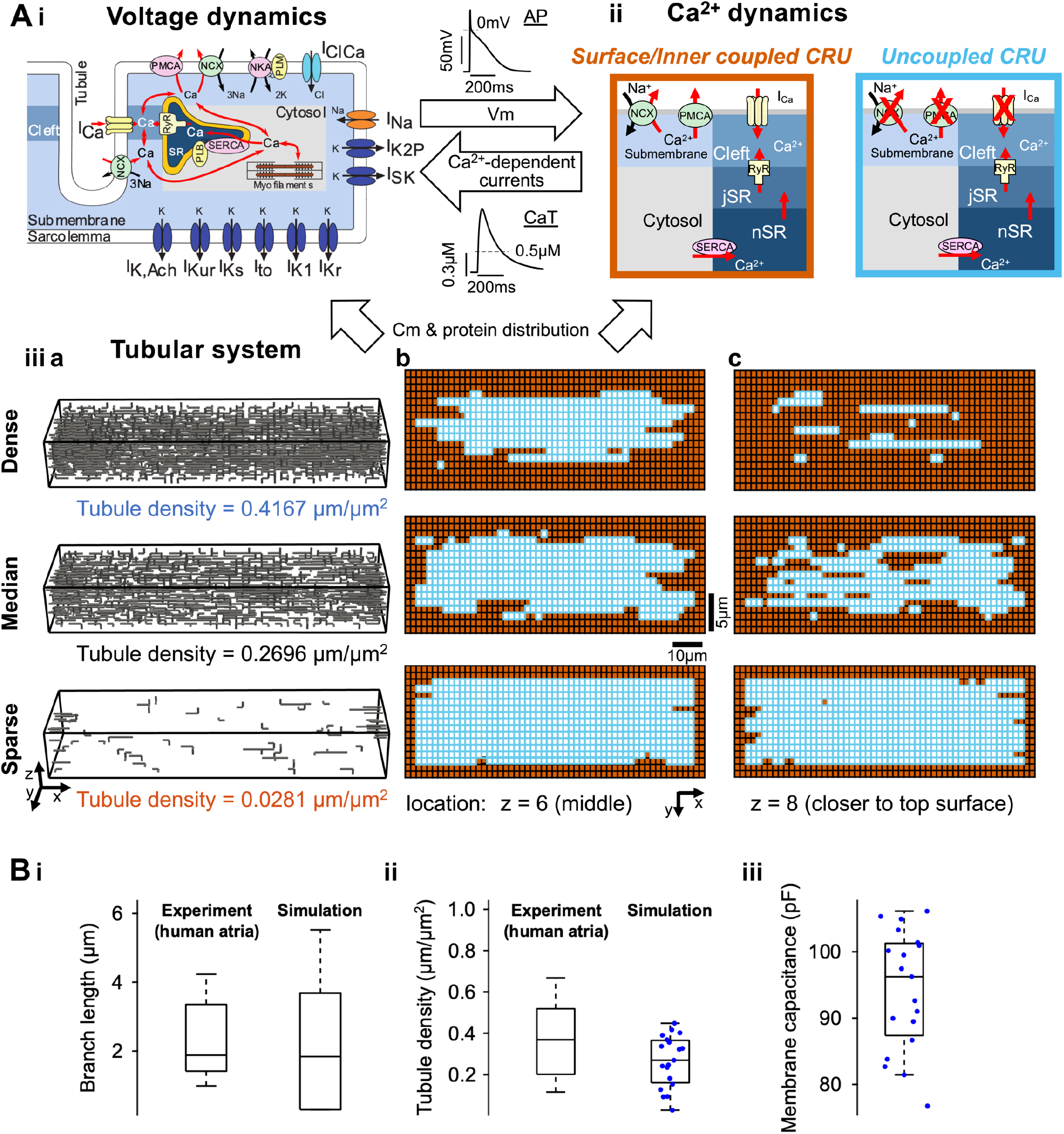
Schematic of model structure and tubular generator features. **A)** Model structure schematic showing membrane voltage dynamics (i) are coupled with Ca^2+^ dynamics (ii) by voltage and Ca^2+^-related currents. Both are also interdependent on the tubular system (iii) whereby tubular structures govern membrane capacitance and sarcolemmal protein distribution. Dependent on ultrastructure, the cell is spatially divided into small Ca^2+^ release units (CRUs). (iiia-iiic) Representative three-dimensional (3D) structures (a) and xy-plane cross-sections of coupled CRUs in central (b) and closer to the top surface (c) of cells with dense (0.4167 μm/μm^2^, top row), median (0.2696 μm/μm^2^, middle row) and sparse (0.0281 μm/μm^2^, bottom row) tubular densities. Due to fewer t-tubules, sparsely tubulated cells have fewer coupled CRUs (orange) in both xy-plane cross-sections. Coupled CRUs are more abundant in upper vs. central xy-planes in both dense and median tubulated cells due to initial tubule formation from the surface sarcolemma or as branches from existing tubules. Each CRU is divided into 5 compartments: cleft, submembrane, cytosol, junctional SR (jSR), and network SR (nSR). Ca^2+^ ions travel (diffuse or get carried) between neighboring compartments and CRUs. Those CRUs coupled with tubules or surface membrane (orange boxes) possess all membrane ion channels and exchangers; the uncoupled CRUs (blue boxes) do not and are devoid of membrane ion channels and exchangers. The membrane currents include LCC Ca^2+^ current (I_Ca_), background sarcolemmal membrane Ca^2+^ current (I_Cabk_), Ca^2+^-activated Cl^-^ current (I_ClCa_), fast Na^+^ current (INa), background Na^+^ current (INabk), small-conductance Ca^2+^-activated K^+^ current (I_SK_), Na^+^/K^+^ pump current (I_Na_K), NCX current (I_NCX_), sarcolemmal membrane Ca^2+^ pump (I_PMCA_), acetylcholine-activated K^+^ current (I_K,Ach_), ultrarapid delayed rectifier K^+^ current (I_Kur_), slowly activating delayed rectifier K^+^ current (I_KS_), transient outward K^+^ current (I_to_), inward rectifier K^+^ current (I_K1_), rapidly activating delayed rectifier K^+^ current (I_Kr_), and 2-pore-domain K^+^ current (I_K2P_). **B)** Estimated tubular branch length (i) and tubular density (ii) in experimental studies (left) and the simulated population (right). The simulated population has 19 randomly generated tubular structures that were used to calculate membrane capacitance (iii). Experimental data are adapted from (Brandenburg et al., 2018).

## Methods

To describe the coupling of electrophysiology, Ca^2+^ signaling and ultrastructure of human atrial cardiomyocytes (**Fig. 1**), we merged the description of surface membrane ion channels/transporters that interact nonlinearly to shape AP dynamics (**Fig. 1Ai**), as described in (Grandi *et al.*, 2011; Morotti *et al.*, 2016a, 2016b), with updated models of the LCC current (I_Ca_) and RyR Ca^2+^ release (I_rel_) (**Fig. 2**) that gate stochastically in individual Ca^2+^ release units (CRUs, **Fig. 1Aii**) (Restrepo *et al.*, 2008; Sato & Bers, 2011). A population of tubular architectures (**Fig. 1Aiii**) was constructed to recapitulate experimental TATS features reported in human atrial myocytes (Brandenburg *et al.*, 2018) and integrated into the model. This ultrastructural detail determines whether any given CRU is coupled with sarcolemmal fluxes or remains uncoupled. The integrative model was reparametrized to recapitulate the rate-dependent properties of electrophysiology and Ca^2+^ dynamics (**Fig. 3A-C**, parameters in **Tables 1-3**). Rigorous model validation was then achieved by testing the capability of the model to simulate properties of electrophysiology, subcellular and whole-cell Ca^2+^ signaling in myocytes with varying TATS structures and subjected to various physiological challenges (**Figs. 2C, 3-6**). Our validated model was then applied to investigate the effects of variable TATS on arrhythmogenesis.

**Figure 2.**
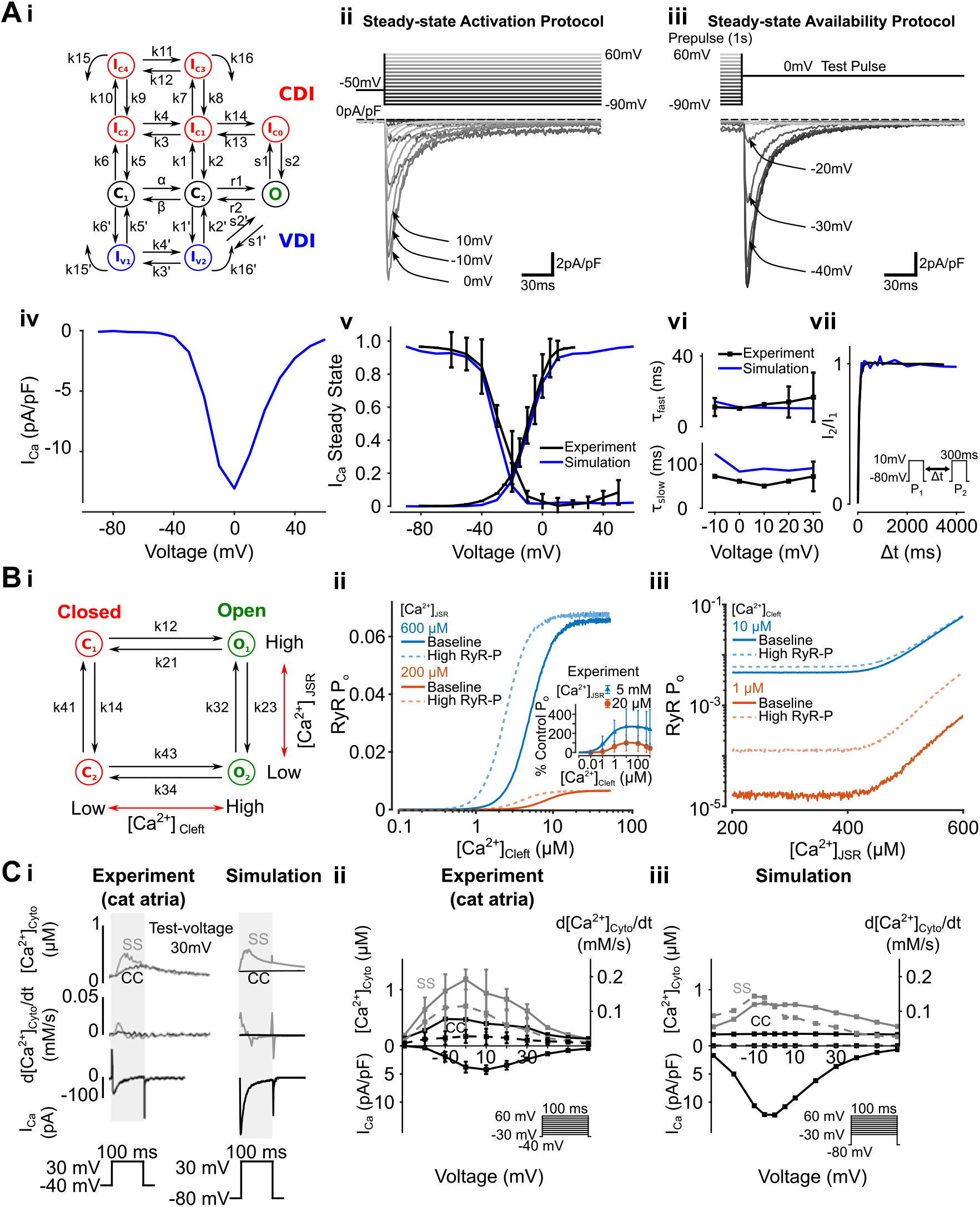
Properties of LCC, RyR, and local Ca^2+^ signaling. **A)** Schematic of LCC Markov model (i), representative traces of I_Ca_ in a cell with median tubules under steady-state activation (ii) and availability (iii) protocols. These data were used to calculate LCC current-voltage (IV) dependence (iv), activation-inactivation curve (v) and fast (top) and slow (bottom) decay time constants (vi). Time-dependent recovery was measured by P1P2 (pulse1-pulse2) protocol (insert), with the ratio of P2- and P1-induced I_Ca_ (I2/I1) recovering steeply to 1.0 with increasing P1-P2 interval (Δt) (vii). Simulation results aligned with experimental observations adapted from (Li and Nattel, 1997). **B)** Schematic of RyR 4-state Markov model (i), dependence of RyR open probability (P_O_) on [Ca^2+^]_cleft_ at [Ca^2+^]_JSR_ of 200 μM and 600 μM (ii), and [Ca^2+^]_JSR_ at [Ca^2+^]_Cleft_ of 1 μM and 10 μM (iii) in a bilayer RyR simulation. Experimental P_O_ dependence on [Ca^2+^]_cleft_ adapted from (Györke and Györke, 1998) showed a similar tendency to the simulated model (ii-inset). RyR hyperphosphorylation (‘High RyR-P’) shifts the dependence of RyR open probability (P_O_) on [Ca^2+^]_cleft_ at [Ca^2+^]_JSR_ of 200 μM and 600 μM (ii), and increased the dependence of P_O_ on [Ca^2+^]_JSR_ at [Ca^2+^]_Cleft_ of 1 μM and 10 μM (iii) in the bilayer RyR simulation. **C)** Simultaneous surface (SS, grey) and cell center (CC, black) traces of subcellular [Ca^2+^]_Cyto_ (top), d[Ca^2+^]_Cyto/dt_ (derivative of [Ca^2+^]_Cyto_ transient, middle) and I_Ca_ (bottom) in an experimental (left) and simulated cell without tubules (right) (i). Voltage dependence of I_Ca_ and local [Ca^2+^]_Cyto_ in experimental (ii) and simulated (iii) cells. Top: local [Ca^2+^]_Cyto_ transient (continuous lines) and d[Ca^2+^]_Cyto/dt_ (dashed lines) in the SS space (grey) and CC region (black). Bottom: voltage dependence of peak I_Ca_ (IV curve). Experimental data represents mean ± S.D., obtained by pacing at multiple test voltages adapted from (Sheehan and Blatter, 2003), with simulated data also obtained using a voltage-clamp stepping protocol. [Na^+^]_Cyto_ was clamped to 0.01 mM for simulation NCX equations to replicate experimental conditions. Simulation results recapitulate the experimental observations that 1) [Ca^2+^]_Cyto_ and d[Ca^2+^]_Cyto/dt_ peaks both occur at the same test-voltage prior to the I_Ca_ peak and 2) the bell-shaped voltage-dependence of [Ca^2+^]_Cyto_ transient amplitude is steeper and increased in amplitude in SS vs. CC.

**Figure 3.**
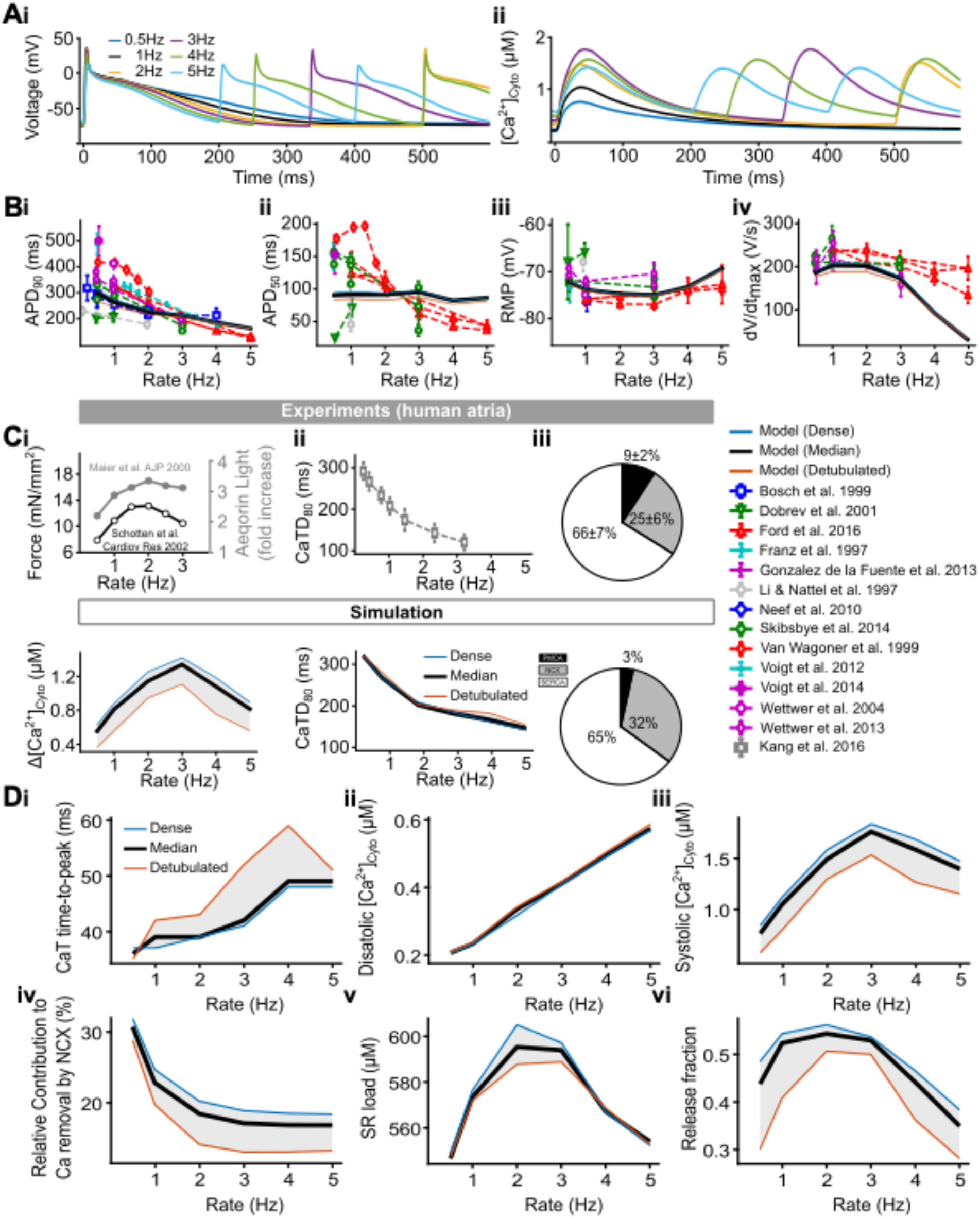
Fitting results for AP and Ca^2+^ transient biomarkers at multiple pacing rates. **A)** Representative AP (i) and Ca^2+^ transient (ii) traces in a cell with median tubules during 0.5, 1, 2, 3, 4, and 5 Hz pacing. **B)** Rate dependent changes in APD_90_ (AP duration at 90% repolarization, i), APD_50_ (ii), RMP (resting membrane potential, iii) and dV/dtmax (maximum upstroke velocity of depolarization, iv) in simulation and experimental observations at 0.5-5 Hz. Models of detubulated (orange), dense (blue) and median (black) tubulated cells are displayed as solid lines with data from previous studies used for fitting referenced in the key below. **C)** Comparison of contractile force/Ca^2+^ transient amplitude (i), CaTD_80_ (Ca^2+^ transient duration at 80% systolic level, ii), and Ca^2+^ extrusion fractions (iii) between experimental results (top) and simulation (bottom). Experimental data of Ca^2+^ extrusion fractions at 0.5 Hz is adapted from (Voigt et al., 2012). **D)** Model prediction of Ca^2+^ signaling rate-dependence properties, including Ca^2+^ transient time to peak (i), diastolic [Ca^2+^]_Cyto_ (ii), systolic [Ca^2+^]_Cyto_ (iii), relative contribution to Ca^2+^ removal by NCX (iv), SR load (v) and release fraction (vi) in detubulated (orange), dense (blue) and median (black) tubulated cells at 0.5-5 Hz stimulation rates.

**Table 1.** Structural parameters for the 3D subcellular model.

| Parameter | Description | Value |
| --- | --- | --- |
| $N$ | Number of CRUs | 10,285 |
| $n_x$ | Number of CRUs along length | 55 |
| $n_y$ | Number of CRUs along width | 17 |
| $n_z$ | Number of CRUs along depth | 11 |
| $length_{\text{CRU}}$ | Length of single CRU | $1.84 \mu\text{m}$ |
| $width_{\text{CRU}}$ | Width of single CRU | $0.9 \mu\text{m}$ |
| $depth_{\text{CRU}}$ | Depth of single CRU | $0.9 \mu\text{m}$ |
| $v_{\text{Cyto}}$ | Local cytosolic volume, CRU | $0.969 \mu\text{m}^3$ |
| $v_s$ | Local submembrane space volume, CRU | $0.025 \mu\text{m}^3$ |
| $v_{\text{JSR}}$ | Local JSR volume, CRU | $0.015 \mu\text{m}^3$ |
| $v_{NSR}$ | Local NSR volume, CRU | 0.025 $\mu\text{m}^3$ |
| $v_{Cleft}$ | Local cleft volume, CRU | 0.126 $\mu\text{m}^3$ |

**Table 2.** Updated Ca^2+^ buffering and time scale of Ca^2+^ diffusion parameters.

| Parameter | Description | Value |
| --- | --- | --- |
| $K_c$ | Dissociation constant of CSQ | 650 μM |
| $K$ | Dimerization constant of CSQ | 900 μM |
| $\tau_{si}$ | Time scale of Ca<sup>2+</sup> diffusion, submembrane to cytoplasm | 0.17 ms |
| $\tau_i^T$ | Time scale of Ca<sup>2+</sup> diffusion, transverse cytosolic | 1.07 ms |
| $\tau_i^L$ | Time scale of Ca<sup>2+</sup> diffusion, longitudinal cytosolic | 1.75 ms |
| $\tau_s^T$ | Time scale of Ca<sup>2+</sup> diffusion, transverse submembrane | 1.28 ms |
| $\tau_s^L$ | Time scale of Ca<sup>2+</sup> diffusion, longitudinal submembrane | 3.06 ms |

**Table 3.**
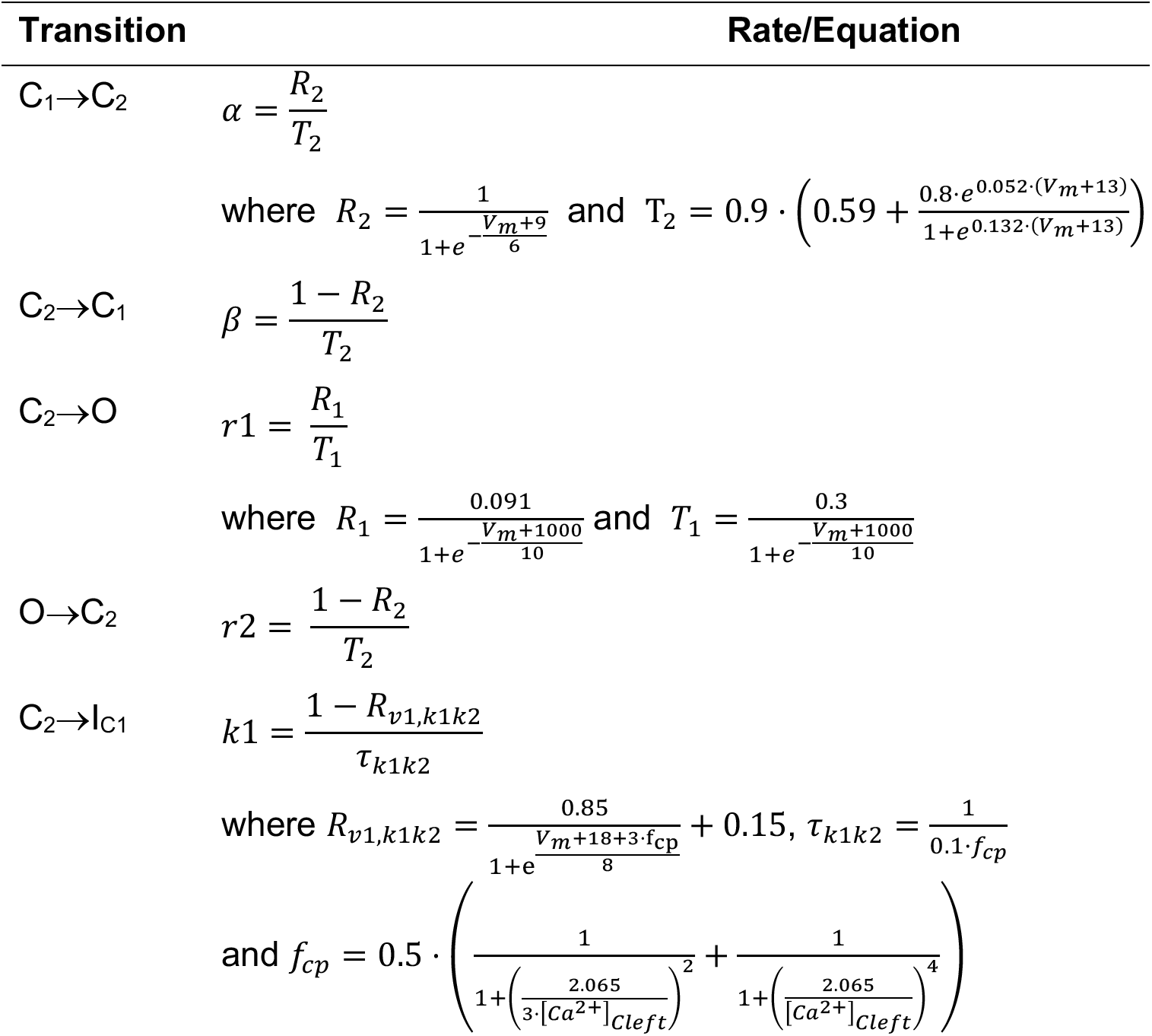
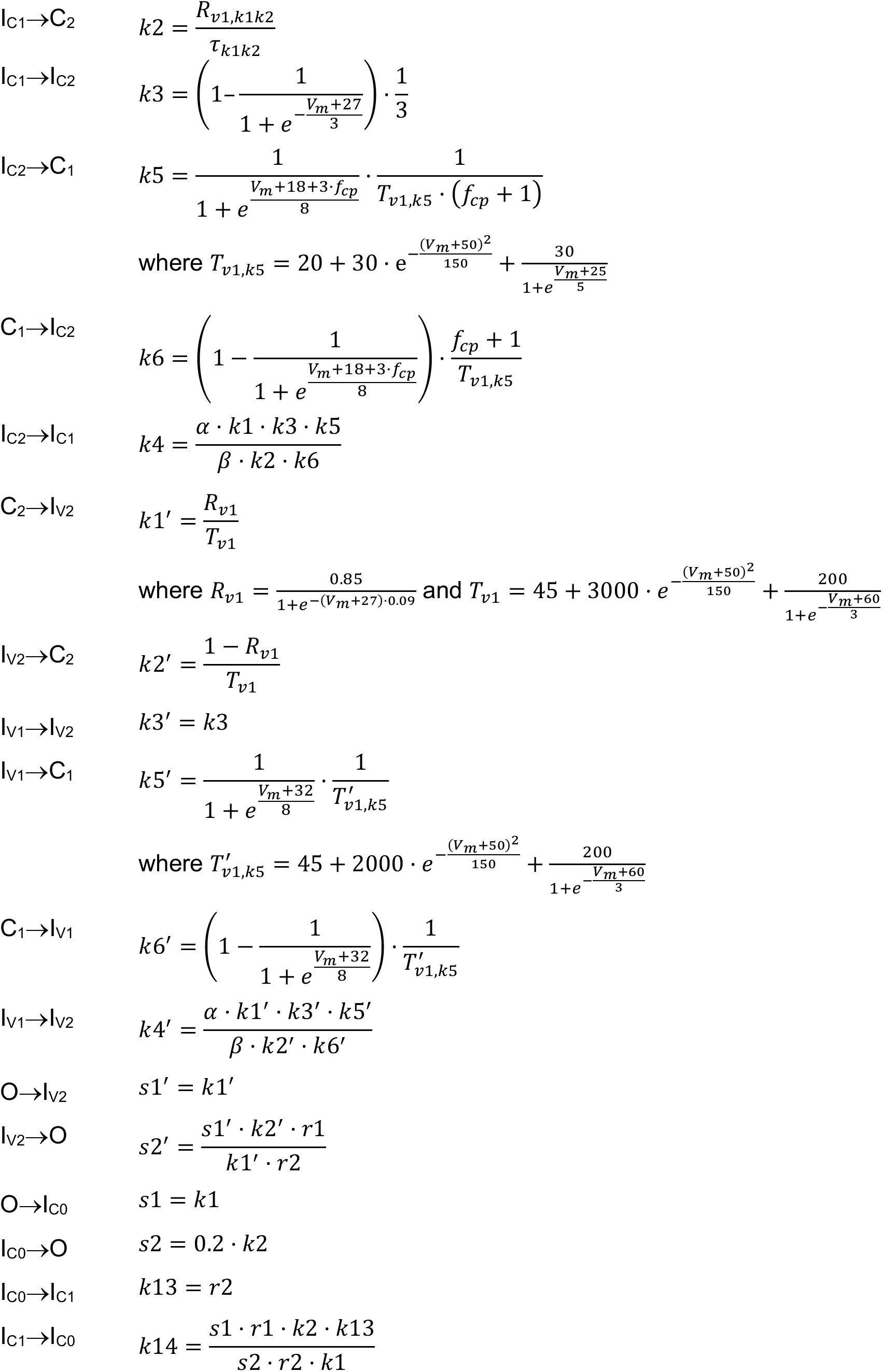
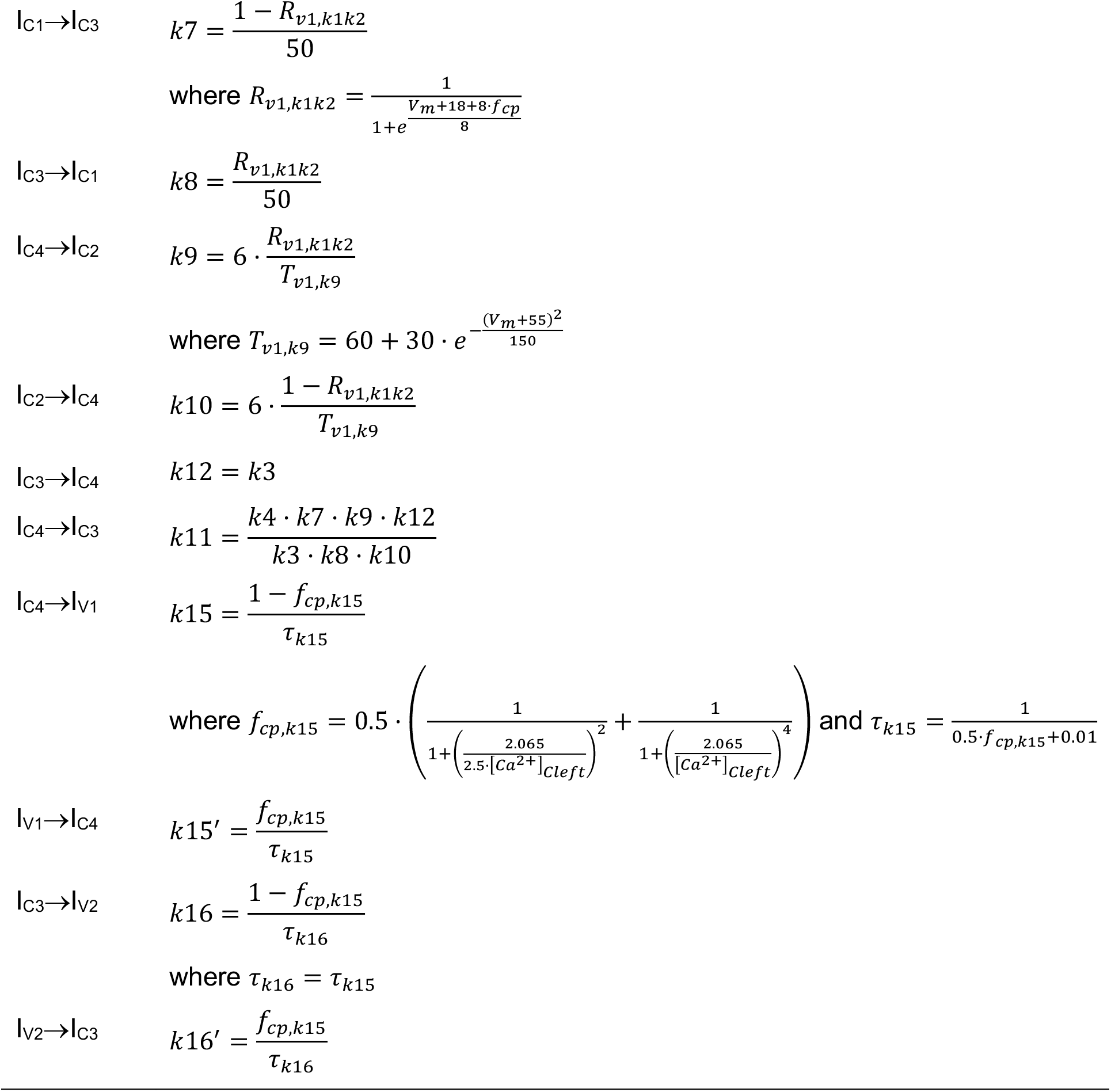
Transition rates of the LCC Markov model.

### Subcellular Ca^2+^ signaling model structure

Our 3D Ca^2+^ signaling model of the human atrial myocyte was based on an established framework for describing rabbit ventricular Ca^2+^ signaling (Restrepo *et al.*, 2008; Sato & Bers, 2011), which accounts for spatiotemporal properties of subcellular Ca^2+^ signaling by simulating a network of CRUs (**Fig. 1Aii**). Here, the cell model dimensions (**Table 1**) and electrical capacitance (**Fig. 1Biii**) were modified to better describe those in human atrial myocytes (Wang Z *et al.*, 1993a, 1993b; Porciatti *et al.*, 1997; Bosch *et al.*, 1999; Polontchouk *et al.*, 2001; Wettwer Erich *et al.*, 2004; Christ T. *et al.*, 2004; Gassanov *et al.*, 2006; Neef *et al.*, 2010; Richards *et al.*, 2011). The new 3D human atrial cell model has a dimension of 101.2 μm × 15.3 μm × 9.9 μm and comprises 10,285 (55 × 17 × 11) CRUs (**Table 1**). Each CRU has five Ca^2+^ compartments, namely the cytosolic, submembrane, and cleft compartments, and the network and junctional sarcoplasmic reticulum (NSR and JSR, respectively), and has a dimension of 1.84 μm × 0.9 μm × 0.9 μm (**Fig. 1Aii**) as in (Sato & Bers, 2011). CRUs are coupled with each other by Ca^2+^ diffusion between respective submembrane, cytosol, and NSR compartments (**Fig. 1Ai-ii**). The cytosolic volume in each CRU was increased to match that in our previous atrial model (Grandi *et al.*, 2011), with the JSR volume decreased to better reproduce the properties of the atrial Ca^2+^ transient (i.e., time-to-peak and amplitude) (**Table 1, Fig. 3Ci, Di**). The cleft volume was the same in each CRU and equal to the average value in (Sato & Bers, 2011). Depending on their position in the cell, we distinguished between peripheral CRUs (i.e., the first two layers close to the cell surface) and inner CRUs (**Fig. 1Aii-iii**). Peripheral and inner CRUs coupled with TATS possess surface membrane ion channels/transporters (i.e., I_Ca_, NCX current, I_NCX_, background Ca^2+^ current, I_Cabk_, plasma membrane Ca^2+^ ATPase (PMCA) current, I_PMCA_, fast Na^+^ current, I_Na_, background Na^+^ current, I_Nabk_, small-conductance Ca^2+^-activated K^+^ current, I_SK_, Ca^2+^-activated Cl^-^ current, I_ClCa_, Na^+^/K^+^ pump current, I_NaK_) distributed in the cleft and/or submembrane space, whereas these processes are absent in the inner CRUs that are not coupled with tubules (inner uncoupled CRUs).

### Generating experiment-based TATS population

A random-walk algorithm as done in (Song *et al.*, 2018) was used to create a population of TATS that quantitatively resemble the features of those reported experimentally in human atrial myocytes (Brandenburg *et al.*, 2018). We converted the experimental TATS features measured in 2D (i.e., total tubular density, branch length, and axial-to-transverse tubule ratio) to 3D features (e.g., we added a z component to the TATS, with z-tubules having the same density as tubules in the transverse direction) and then assigned them as inputs to our random-walk-based algorithm. The constructed 3D TATS networks were subsequently verified by extracting parameters in 2D cross-sections and confirming that the TATS properties matched experimental measurements (**Fig. 1Bi-ii**). Our simulated population contains 20 TATS configurations with tubular density ranging from fully detubulated to that of densely tubulated atrial cardiomyocytes. Representative simulated dense/median/sparse TATS are in **Fig. 1Aiii**. TATS density is greater (i.e., a larger number of coupled CRUs are present) near the cell surface than in the cell center as tubules invaginate from the cell surface and/or branch from existing tubules (Richards *et al.*, 2011).

The coupling of TATS with voltage dynamics and subcellular Ca^2+^ signaling dictated which CRUs in the network are coupled with the sarcolemma (**Fig. 1Aii-iii**), and affected the membrane electrical capacitance as follows:

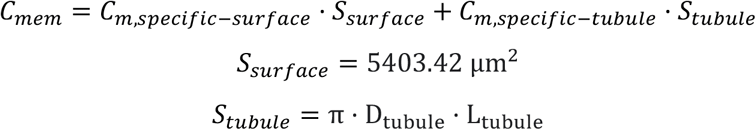

where we assume a larger specific capacitance of the surface sarcolemma *C_m,specific–surface_* = 0.01 *pF* · *μm*^-2^ compared to the specific capacitance of tubular sarcolemma *C_m,specific–tubule_* = 0.0056 *pF* · *μm*^-2^, as estimated in experimental measurements (Pásek *et al.*, 2008); D_tubule_ = 0.3 μm is the tubule diameter, as estimated in (Brandenburg *et al.*, 2018), and L_tubule_ is the total length of tubules, which is associated with TATS density and varies with each TATS configuration. The median *C_mem_* of our cell population is 96.25 pF (**Fig. 1Biii**), which is similar to experimental estimates in human atrial myocytes (Van Wagoner *et al.*, 1999; Christ T. *et al.*, 2004; González de la Fuente *et al.*, 2013; Brandenburg *et al.*, 2018).

### Ca^2+^ buffering and diffusion

Descriptions of Ca^2+^ buffering in the cleft, submembrane, and cytosolic compartments were replaced with the kinetic binding schemes in our previous models (Grandi *et al.*, 2011; Morotti *et al.*, 2016b). Ca^2+^ buffering by calsequestrin (CSQ) in the JSR was modeled using the rapid equilibrium approximation as done previously (Restrepo *et al.*, 2008; Sato & Bers, 2011). We updated the CSQ dissociation constant (*K_C_*) based on (Grandi *et al.*, 2011), and modified CSQ dimerization constant (*K*) to an intermediate value between those used in (Restrepo *et al.*, 2008) and (Sato & Bers, 2011), to match experimental RyR-[Ca^2+^]_cleft_ dependence (**Fig. 2Bii insert**) (Györke & Györke, 1998). Updated parameters are listed in **Table 2**.

The diffusion equations as in (Sato & Bers, 2011) were used with updated time scales of Ca^2+^ diffusion (**Table 2**). Specifically, we slowed the diffusion between submembrane and cytosol to prolong the time to peak of the Ca^2+^ transient, and hastened the Ca^2+^ diffusion among neighboring submembrane spaces to promote Ca^2+^ diffusion between neighboring CRUs. In addition, diffusion among neighboring cytosolic compartments was accelerated based on multiple experimental observations (Wang, 1953; Kushmerick & Podolsky, 1969; Donahue & Abercrombie, 1987; Allbritton *et al.*, 1992; Cordeiro *et al.*, 2001; Wu & Bers, 2006; Picht *et al.*, 2011) and experiment-based evaluations (Swietach *et al.*, 2010; Bers & Shannon, 2013).

### Dynamics of Ca^2+^ cycling in CRU

The Ca^2+^ concentration in the Ca^2+^ compartments of each CRU was calculated by accounting for all transmembrane (sarcolemmal and SR) Ca^2+^ currents and fluxes, Ca^2+^ buffers, and Ca^2+^ diffusion between neighboring compartments:

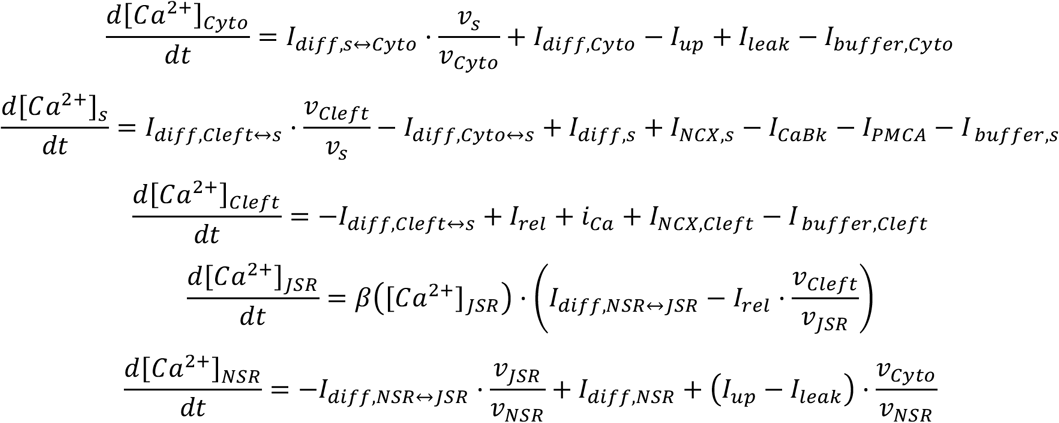

where [*Ca*^2+^]_*x*_ is the local Ca^2+^ concentration in cytosolic (x: Cyto), submembrane (x: s), cleft (x: Cleft), NSR (x: NSR), or JSR (x: JSR) compartments; *I_diff,x_* is the diffusive current between neighboring compartments; and *I_up_* is the SERCA uptake current; *I_leak_* is the SR leak current; *β*([*Ca*^2+^]_*JSR*_) is the rapid equilibrium approximation term of the CSQ Ca^2+^ buffering for [*Ca*^2+^]_*JSR*_; *I_buffer,x_* is the dynamic buffer current in compartments, described as follows:

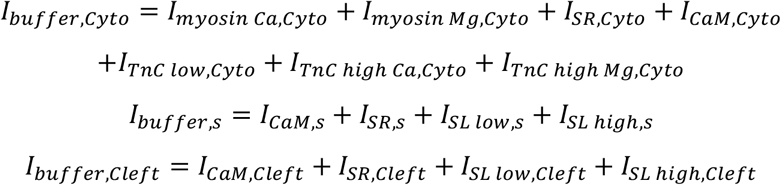

### Updated ion channels and transporters

An updated version of the Grandi et al. human atrial myocyte model (Grandi *et al.*, 2011) was used as described in (Morotti *et al.*, 2016a, 2016b) (**Fig. 1Ai**). A new Markov model of the LCC current was developed and parameters describing RyR release and SR Ca^2+^-ATPase (SERCA) uptake were modified, as described below, to recapitulate characteristics of Ca^2+^ signaling (**Fig. 2C and 3C**). We further added descriptions of small-conductance Ca^2+^-activated K^+^ current (I_SK_) as in (Morotti *et al.*, 2016a), and replaced the model formulations of I_Na_ and I_NCX_ with the INa model from (Courtemanche *et al.*, 1998) and the I_NCX_ model from (Soltis & Saucerman, 2010). The maximal conductance and transporter rates of I_SK_, I_NCX_, I_CaBk_, I_CaP_, I_NaK_, I_Na_, and I_NaBk_ were adjusted to better reproduce the rate-dependence of AP and Ca^2+^ dynamics.

All membrane ion channels and transporters are distributed uniformly to all coupled CRUs. Within each coupled CRU, the LCCs are assumed to be in the cleft area and are closely coupled with RyRs as in (Restrepo *et al.*, 2008; Sato & Bers, 2011). NCXs are distributed with 11% of the proteins in the cleft and 89% in the submembrane as (Grandi *et al.*, 2011), similar to I_NaBk_, I_NaK_, I_Na_, I_ClCa_, I_SK_, while I_Cabk_ and I_PMCA_ are located only in the submembrane as in (Sato & Bers, 2011).

#### I_Ca_ Markov model

Each CRU contains 4 LCCs, as in (Restrepo *et al.*, 2008). We developed a novel 10-state stochastic Markov model (**Fig. 2Ai**) based on (Morotti *et al.*, 2012) to replicate biophysical properties of I_Ca_ in human atria and recapitulate two distinct components of inactivation (Li & Nattel, 1997). This model has 1 open state (O), 2 closed states (C_1_, C_2_), 5 Ca^2+^-dependent inactivation states (I_C0_, I_C1_, I_C2_, I_C3_, I_C4_), and 2 voltage (V_m_)-dependent inactivation states (I_V1_, I_V2_). The transition rates of the Markovian *I_Ca_* current are described in **Table 3**.

The model tracks the total number of LCCs residing in the open state, with whole-cell I_Ca_ calculated by the sum of unitary current (*i_Ca_*), as described using the original model formulation in (Sato & Bers, 2011). We added descriptions of Na^+^ and K^+^ fluxes through LCCs from (Grandi *et al.*, 2011; Morotti *et al.*, 2016b). To calculate the unitary Na^+^ and K^+^ currents (*i_Ca,Na_* and *i_Ca,K_*), the Na^+^ and K^+^ permeability in the original model was divided by the total number of LCCs (**Table 4**).

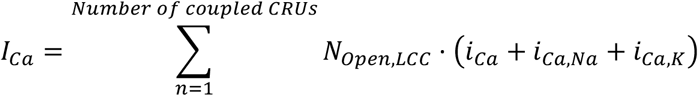

Our new I_Ca_ model reproduced steady-state activation and inactivation curves (**Fig. 2Aii, iiii, and v**), the fast and slow inactivation time constants (**Fig. 2Avi**), and time-dependent recovery (**Fig. 2Avii**), which resembles the experimental measurements (Li & Nattel, 1997).

**Table 4.** Updated human atrial myocyte ionic parameters.

| Parameter | Description | Value |
| --- | --- | --- |
| $P_{Ca}$ | Unitary LCC $Ca^{2+}$ permeability ($I_{Ca}$) | $8.19 \text{ } \mu\text{mol} \cdot \text{C}^{-1} \cdot \text{ms}^{-1}$ |
| $P_{Na}$ | Unitary LCC $Na^{+}$ permeability ($I_{Ca}$) | $6.14 \cdot 10^{-13} \text{ cm} \cdot \text{s}^{-1}$ |
| $P_K$ | Unitary LCC $K^{+}$ permeability ($I_{Ca}$) | $1.1 \cdot 10^{-11} \text{ cm} \cdot \text{s}^{-1}$ |
| $g_{SK}$ | Small-conductance $Ca^{2+}$-activated<br>$K^{+}$ conductance per CRU ($I_{SK}$) | $2.21 \cdot 10^{-10} \text{ mS}$ |
| $k_{d,SK}$ | Dissociation constant of intracellular $Ca^{2+}$ ($I_{SK}$) | $0.79 \cdot 10^{-3.45} \text{ mM}$ |
| $N_{RyR}$ | Number of RyRs per CRU ($I_{rel}$) | 41 |
| $J_{max}$ | Strength of $Ca^{2+}$ release flux ($I_{rel}$) | $0.1321 \text{ } \mu\text{m}^3 \cdot \text{ms}^{-1}$ |
| $K_{cp}$ | Constant ($I_{rel}$) | 10 $\mu\text{M}$ |
| $h_{cp}$ | $[Ca^{2+}]_{cleft}$-dependence exponent of $k_{12}$ and $k_{43}$ ($I_{rel}$) | 3 |
| $v_{NCX}$ | Strength of the NCX $Ca^{2+}$ flux per CRU ($I_{NCX}$) | $17.88 \text{ } \mu\text{M} \cdot \text{ms}^{-1}$ |
| $G_{ClCa}$ | $Ca^{2+}$-dependent $Cl^{-}$ conductance per CRU ($I_{ClCa}$) | $6.5 \cdot 10^{-10} \text{ mS}$ |
| $G_{CaBk}$ | Strength of the background SL $Ca^{2+}$ flux per CRU ($I_{CaBk}$) | $0.0014 \text{ } \mu\text{M} \cdot \text{ms}^{-1}$ |
| $V_{max}$ | Strength of the PMCA current per CRU ($I_{PMCA}$) | $0.0097 \text{ } \mu\text{M} \cdot \text{ms}^{-1}$ |
| $I_{bar,NaK}$ | $Na^{+}/K^{+}$ pump maximal transport rate per CRU, ($I_{NaK}$) | $1.49 \cdot 10^{-14} \text{ A}$ |
| $g_{Na}$ | $Na^{+}$ conductance per CRU ($I_{Na}$) | $1.07 \cdot 10^{-7} \text{ mS}$ |
| $g_{NaB}$ | $Na^{+}$ background conductance per CRU ($I_{NaBk}$) | $7.09 \cdot 10^{-12} \text{ mS}$ |
| $v_{up}$ | Strength of uptake per CRU ($I_{up}$) | $0.36 \text{ } \mu\text{M} \cdot \text{ms}^{-1}$ |
| $K_i$ | $K_{mf}$ for SR Ca pump forward mode ($I_{up}$) | 0.615 $\mu\text{M}$ |

#### SR Ca^2+^ release and uptake

The 4-state stochastic RyR Markov model (**Fig. 2Bi**) in (Restrepo *et al.*, 2008; Sato & Bers, 2011) was modified to incorporate human atrial myocyte-specific characteristics. This model includes 2 open states (O_1_, O_2_) and 2 closed states (C_1_, C_2_), whereby transitions are regulated by [Ca^2+^]_Cleft_ and [Ca^2+^]_JSR_. The RyR number per CRU (NRyR) was reduced to 41 based on the experimental measurement (Boyd *et al.*, 2018) and the strength of SR Ca^2+^ release flux (J_max_) was reduced compared to (Sato & Bers, 2011) to match the rate-dependence of Ca^2+^ transient amplitude and duration (**Fig. 3Ci-ii**). Furthermore, we left shifted the [Ca^2+^]_cleft_-dependence (Kcp) and increased the hill coefficient (hcp) (**Table 4**) to match the [Ca^2+^]_Cleft_-dependence of the RyR P_O_ (Györke & Györke, 1998; Fill & Gillespie, 2018). Our RyR model reproduces the changes in RyR P_O_ when [Ca^2+^]_Cleft_ or [Ca^2+^]_JSR_ are varied (**Fig. 2Bii-iii**), similar to the observations from lipid-bilayer RyR experiments (**Fig. 2Bii, insert**) (Györke & Györke, 1998). Finally, *k*_32_ was updated to impose reversibility of the Markov model. RyR hyperphosphorylation was simulated by left shifting the [Ca^2+^]_cleft_-dependence of SR release (i.e., reducing K_cp_ by 50%) to increase RyR P_O_ (**Fig. 2Bii-iii**). The transition rates of the RyR Markov model are described in **Table 5**.

**Table 5.** Transition rates of the RyR Markov model.

| Transition | Rate/Equation |
| --- | --- |
| $C_1 \rightarrow O_1$ | $k_{12} = K_u \cdot \frac{([Ca^{2+}]_{cleft})^{h_{cp}}}{K_{cp}^{h_{cp}} + ([Ca^{2+}]_{cleft})^{h_{cp}}} + w$<br>where $K_{cp} = 10 \mu M$, $h_{cp} = 3$, $K_u = 5 ms^{-1}$ and $w = 0.000001 ms^{-1}$ |
| $C_2 \rightarrow O_2$ | $k_{43} = K_b \cdot \frac{([Ca^{2+}]_{cleft})^{h_{cp}}}{K_{cp}^{h_{cp}} + ([Ca^{2+}]_{cleft})^{h_{cp}}} + w$<br>where $K_b = 0.005 ms^{-1}$ |
| $O_1 \rightarrow C_1$ | $k_{21} = 0.5 ms^{-1}$ |
| $C_2 \rightarrow O_2$ | $k_{34} = 3.3 ms^{-1}$ |
| $C_2 \rightarrow C_1$ | $k_{41} = 0.008 ms^{-1}$ |
| $C_1 \rightarrow C_2$ | $k_{14} = \frac{\hat{M}([Ca^{2+}]_{JSR}) \cdot \tau_b^{-1} \cdot B_{CSQ}}{B_{CSQ}^0}$<br>where $\hat{M}([Ca^{2+}]_{JSR}) = \frac{\sqrt{1+8 \cdot \rho \cdot B_{CSQ}}-1}{4 \cdot \rho \cdot B_{CSQ}}$, $\rho = \frac{5000}{1 + \left(\frac{K}{[Ca^{2+}]_{JSR}}\right)^{23}}$<br>$\tau_b = 0.5 ms$, $B_{CSQ} = 400 \mu M$, $B_{CSQ}^0 = 400 \mu M$, and $K = 900 \mu M$ |
| $O_1 \rightarrow O_2$ | $k_{23} = k_{14}$ |
| $O_2 \rightarrow O_1$ | $k_{32} = \frac{k_{41} \cdot k_{12} \cdot k_{23} \cdot k_{34}}{k_{43} \cdot k_{21} \cdot k_{14}}$ |

SERCA uptake current (*I_up_*) was modified from (Sato & Bers, 2011) by reducing the Ca^2+^ affinity (increasing *K_i_*) as in (Grandi *et al.*, 2011) and adjusting the maximum Ca^2+^ uptake rate *v_up_* to reproduce the rate-dependence of Ca^2+^ transient duration (**Fig. 3Cii**) (Kang *et al.*, 2016) and the relative contribution of SERCA to Ca^2+^ removal during the Ca^2+^ transient (**Fig. 3Ciii**) (Voigt *et al.*, 2012) seen experimentally. **Table 4** contains a list of the updated parameters.

#### Small-conductance Ca^2+^-activated K^+^ current (I_SK_)

I_SK_ was added to our 3D subcellular model from (Morotti *et al.*, 2016a) with adjusted current maximal conductance (*g_SK_*),

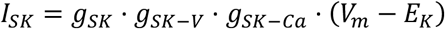

where V_m_- and Ca^2+^-dependent components are formulated as

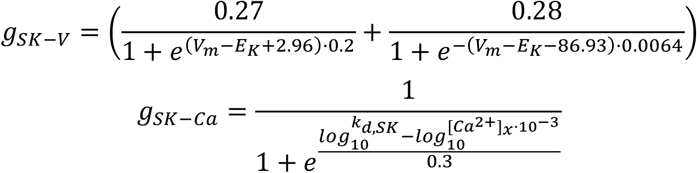

and [*Ca*^2+^]_*x*_ is Ca^2+^ concentration in the cleft or submembrane space and *E_K_* is the Nernst potential for K^+^.

#### Other updated currents

Maximum current strength/flux parameters of I_NCX_ (Soltis & Saucerman, 2010), I_ClCa_ (Morotti *et al.*, 2016b), I_Cabk_ (Sato & Bers, 2011), I_PMCA_ (Sato & Bers, 2011), I_NaK_ (Morotti *et al.*, 2016b), I_Na_ (Courtemanche *et al.*, 1998), I_Nabk_ (Grandi *et al.*, 2011) were adjusted (**Table 4**) to recapitulate the rate-dependent dynamic properties of AP and Ca^2+^ (**Fig. 3A-C**).

### Numerical Method

Our new 3D human atrial cell model was implemented in C++ and parallelized using OpenMP 5.1 (Dagum & Menon, 1998). The ordinary differential equations (ODEs) and subcellular Ca^2+^ diffusion were solved using an explicit Forward Euler method except that I_Ca_ and RyR gating behaviors were described stochastically as in (Restrepo *et al.*, 2008; Sato & Bers, 2011), and that ODEs of I_Na_ were solved using the Rush-Larsen scheme (Rush & Larsen, 1978). The integration time step was 0.01 ms.

### Simulation protocols

To test the rate dependence of AP and Ca^2+^ dynamics in human atrial myocytes, a pacing-and-pause protocol was applied. The protocol starts with a 4-sec unstimulated period, followed by a 28-sec long stimulation period, in which the myocyte is paced at 0.5, 1, 2, 3, 4, or 5 Hz by injecting a depolarizing current (5 ms in duration and 12.5 pA/pF in strength), and ends with a 5-sec pause. In all simulations, the model state variables were assigned with the same initial conditions. While our model does not describe dynamic Na^+^ handling, which slowly varies (within minutes) with the stimulation frequency, we incorporated a rate-dependent description of Na^+^, i.e., the intracellular Na^+^ concentration ([Na^+^]_Cyto_, [mM]) is described as a function of the basic cycle length (BCL, [ms]) (Restrepo *et al.*, 2008; Sato & Bers, 2011) to match the Na^+^-BCL relation simulated using (Grandi *et al.*, 2011). Using rate-corrected [Na^+^]_Cyto_ values allows the virtual cell to rapidly reach steady-state when the pacing frequency is varied.

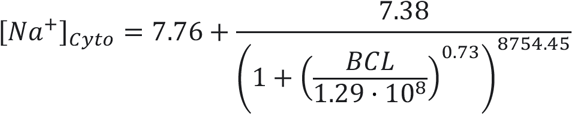

LCCs and RyRs gate stochastically in our model. To ensure the robustness of our main simulations results and conclusions, we simulated each cell with 10 different random seeds.

### Detection and analysis of Ca^2+^ sparks, spontaneous Ca^2+^ release events (SCRs) and delayed after-depolarizations (DADs)

The [Ca^2+^]_Cyto_ waveform starting at the peak of the last paced beat with the subsequent no-stimulation period was used for analysis. Spark and SCR peaks were identified in MATLAB R2021 utilizing the *findpeak* function, with a ‘MinPeakProminence’ of 0.05 μM and ‘MinPeakDistance’ of 100 ms used to define sparks. Amplitudes of SCR events were calculated as the difference between the peak and the minimum [Ca^2+^]_Cyto_ values. To determine the properties of SCR, a detection threshold of 0.3 μM was used as the minimum SCR amplitude. Likewise, membrane voltage waveforms were also examined in MATLAB, with DADs detected when the membrane depolarization amplitude was greater than 10 mV (**Fig. 7A**). Rate threshold of SCRs and DADs was determined as the lowest pacing rate that produced the respective arrhythmic events. To study local Ca^2+^ changes, we averaged the [Ca^2+^]_Cyto_ within surface CRUs, inner coupled CRUs (those CRUs located away from the cell surface but coupled to tubules), and inner uncoupled CRUs.

### Source code

The source code of our new 3D human atrial cell model can be accessed from http://elegrandi.wixsite.com/grandilab/downloads and https://github.com/drgrandilab.

## Results

### Fitting of AP and Ca^2+^ transient biomarkers in the myocyte model against experimental observations in the human atria

To investigate the effects of varying TATS on whole-cell and subcellular Ca^2+^ signaling and electrophysiology in the human atrial myocyte, we built a mathematical model coupling voltage, spatially-detailed Ca^2+^ dynamics, and TATS (**Fig. 1A**) that integrates a vast array of experimental data from various published sources. A detailed description of the ultrastructural characteristics (**Fig. 1B, Tables 1-2**), ion channel and transporter properties (**Fig. 2A-B, Tables 3-5**), and subcellular Ca^2+^ handling features (**Fig. 2C**) in human atrial myocytes used to parameterize our model are provided in the Methods. Given the model complexity and large number of parameters, we also included human atrial electrophysiologic and global Ca^2+^-handling biomarkers (**Fig. 3**) in the fitting process. The representative traces of steady-state APs and Ca^2+^ transients at varying pacing rates (**Fig. 3A**) and simulated properties indicate that fast pacing varied AP and Ca^2+^ transient biomarkers in agreement with experimental data from multiple sources (**Fig. 3B-C**). The model was mainly fitted to mimic the rate-dependence of the AP duration (APD, **Fig. 3Bi**), Ca^2+^ transient amplitude (**Fig. 3Ci**), and Ca^2+^ transient duration (**Fig. 3Cii**). Experimental observations of relative contributions to Ca^2+^ removal by PMCA, NCX, and SERCA were also replicated by our model (**Fig. 3Ciii**). While we utilized the model with median tubular structure (black solid lines), the AP properties above were not markedly changed when simulating models with less or more dense TATS (orange and blue solid lines, respectively). However, lack of TATS was associated with reduced Ca^2+^ transient amplitude (**Fig. 3Ci**) resulting from reduced systolic Ca^2+^ (**Fig. 3Diii**) and fractional release (**Fig. 3Dvi**) and longer time to peak (**Fig. 3Di**). In addition, the biphasic dependence of Ca^2+^ transient amplitude on the pacing rate results from analogous dependence on systolic Ca^2+^ levels (**Fig. 3Diii**), SR Ca^2+^ content (**Fig. 3Dv**), and fractional release (**Fig. 3Dvi**). Overall, these data indicate that our model recapitulates the main electrophysiologic and whole-cell Ca^2+^ handling properties in human-atria-specific experiments and also their rate dependence.

### Validation of APs, subcellular and global Ca^2+^ transient and TATS biomarkers in the myocyte model against experimental observations in the human atria

Next, we validated the model to verify its ability to predict an independent experimental dataset. We focused on AP properties, subcellular Ca^2+^ spark and wave characteristics, and global Ca^2+^ transient features measured when challenging atrial myocytes with ion channel blockers or osmotic shock (to disrupt TATS).

As shown in experiments (Van Wagoner *et al.*, 1999), simulating the application of the I_Ca_ blocker nifedipine shortened the APD and weakened the APD rate dependence (**Fig. 4B** vs. **Fig. 4A**). The summary data shows that the percentage changes in APD90 (with respect to the value at 2 Hz pacing) are similar in the experimental (**Fig. 4Ci**, adapted from (Van Wagoner *et al.*, 1999) and simulation results (**Fig. 4Cii**). Thus, the simulated APD rate-dependence replicates the response to LCC blockade measured in human atrial myocytes.

**Figure 4.**
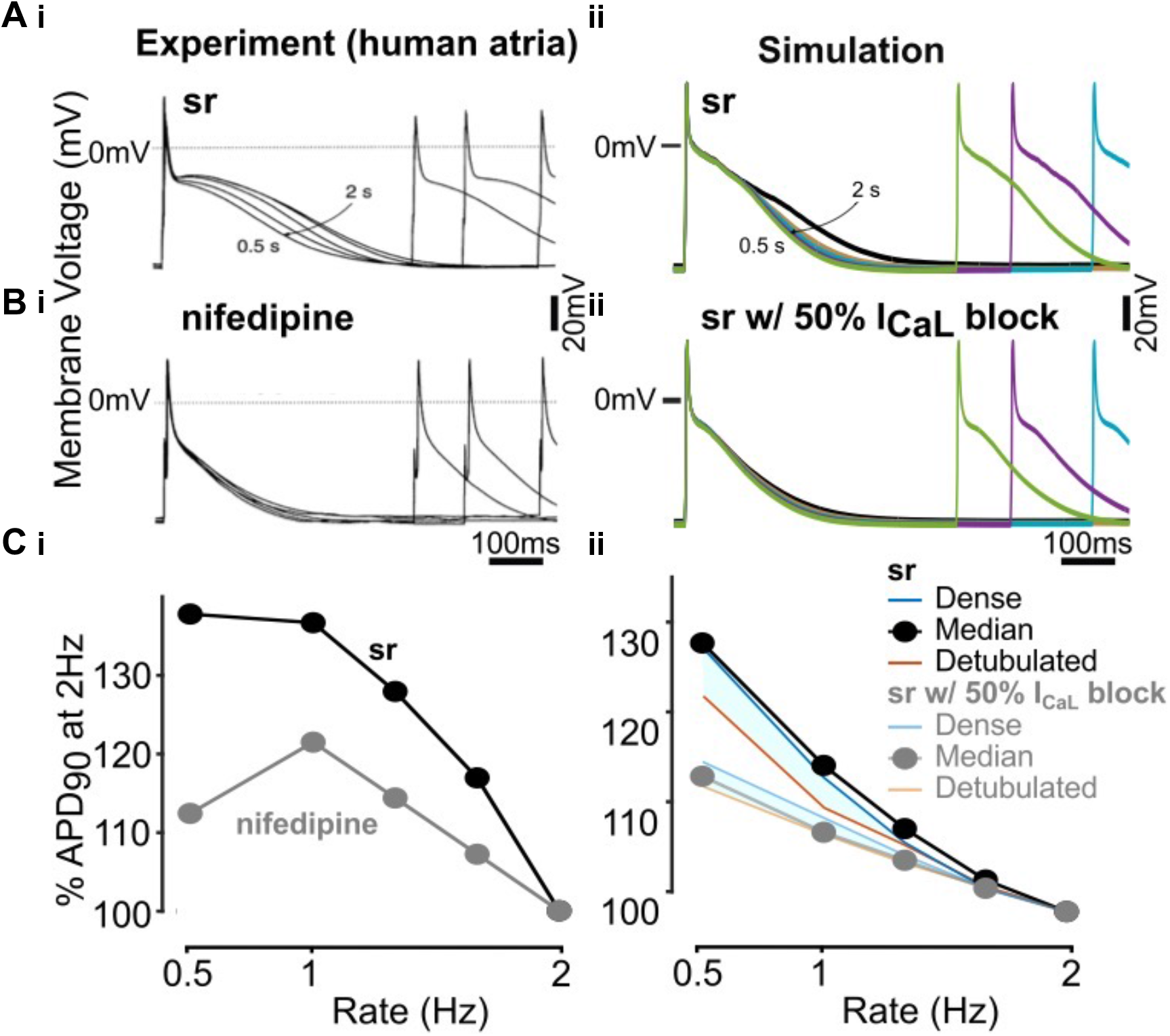
Validation of electrophysiology by I_Ca_ block with and without nifedipine. **A)** Representative AP traces showing AP rate-dependence in normal sinus rhythm (sr) in experimental (Van Wagoner David R. et al., 1999) (i) and simulated (ii) cells. Simulated cells with representative tubular structures (Dense, Median, and Detubulated) were paced with the basic cycle length of 2, 1, 0.75, 0.6, and 0.5 s. **B)** Representative AP traces showing AP rate-dependence with I_Ca_ block by nifedipine or 50% I_Ca_ reduction in experimental (i) and simulated (ii) cells. In the simulation, L-type Ca^2+^ channels were blocked by 50% in the common pool model (Grandi et al., 2011), then paced at the same pacing rates to steady-state. The values of steady-state [Na^+^]_Cyto_ in the common pool model were used to update the [Na^+^]_Cyto_ equations in the 3D model prior to blocking L-type Ca^2+^ channels by 50% in the 3D model during pacing.

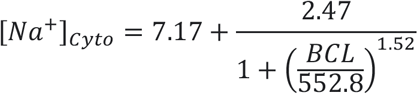 **C)** Summary data showing percentage change in APD_90_ at varying stimulation rates is similar in the experiment (i) and simulation (ii) results.

To validate the predicted subcellular and whole-cell properties of Ca^2+^ signaling, we analyzed the spatiotemporal characteristics of the intracellular Ca^2+^ transients evoked by electrical stimulation and caffeine application. The simulated transversal linescan images show AP-induced Ca^2+^ waves propagating from the cell periphery to the cell center as shown in human atrial myocyte experiments (**Fig. 5A**) (Greiser *et al.*, 2014). The caffeine-evoked Ca^2+^ transient was larger and the release was more synchronous than the electrically evoked transient (**Fig. 5B**). Notably, the ratio of central to surface Ca^2+^ transient amplitude (cc/ss (ratio)) was similar between experiments in rabbit atrial myocytes (Greiser *et al.*, 2014) and human atrial myocyte simulation. Our model also recapitulates the regional differences in the surface vs. central [Ca^2+^]_Cyto_ and d[Ca^2+^]_Cyto/dt_ detected in voltage-clamp experiments in cat atrial myocytes that lack TATS, whereby [Ca^2+^]_Cyto_ and d[Ca^2+^]_Cyto/dt_ are larger at the cell surface, the bell-shaped voltage-dependence of the [Ca^2+^]_Cyto_ transient amplitude is steeper, and [Ca^2+^]_Cyto_ and d[Ca^2+^]_Cyto/dt_ peak at the same test-voltage prior to the I_Ca_ peak (**Fig. 2C**) (Sheehan & Blatter, 2003). To evaluate whole-cell Ca^2+^ signaling, we compared the simulated Ca^2+^ transient characteristics with measurements in human atrial myocytes and found good agreement in the predicted vs. measured (Voigt *et al.*, 2012, 2014) diastolic and systolic twitch Ca^2+^ concentration and in the amplitude and decay time constant of the caffeine-induced Ca^2+^ transient (**Fig. 5C**). Overall, simulated properties of Ca^2+^ transients and waves match experimental observations in human and rabbit atrial myocytes.

**Figure 5.**
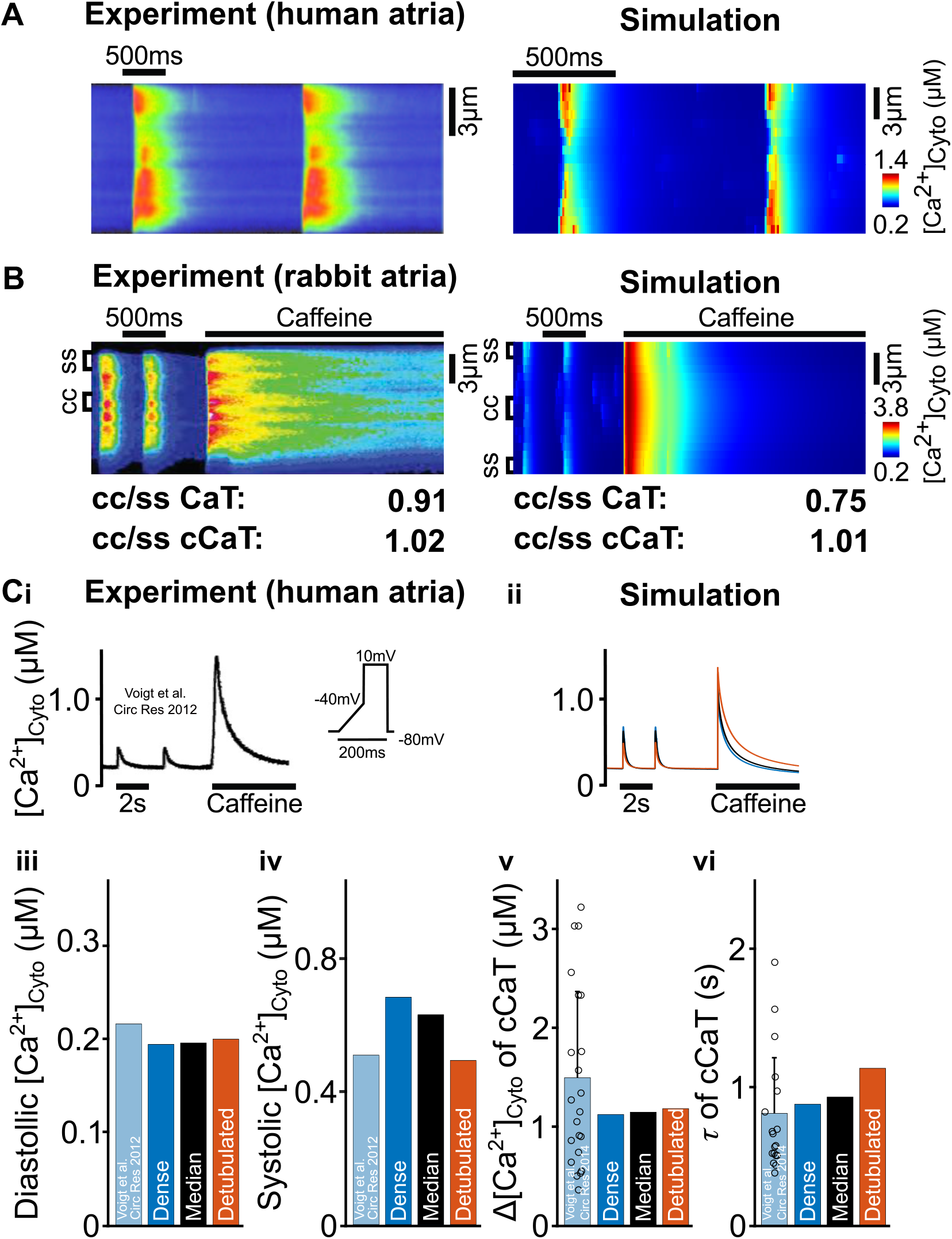
Validation of local and global Ca^2+^ signaling with and without caffeine-evoked SR depletion. **A)** Linescan images showing Ca^2+^ release in the experimental (Greiser et al., 2014) (0.5 Hz, left) and simulated median tubulated (1Hz, right) atrial cells. The simulated cell was paced for 28 seconds to reach steady-state, and the local [Ca^2+^]_Cyto_ of 17 CRUs along the central y-axis during the last 2 beats was measured to get the spatiotemporal image. **B)** Linescan images showing SR Ca^2+^ release at the cell surface and in the center following respective caffeine application or clamping RyR P_O_ at 0.2 in experimental (Greiser et al., 2014) (left) and simulated median tubulated (right) cells. The simulated cell was paced at 0.5 Hz, with caffeine application 750 ms after the last stimuli generating SR Ca^2+^ release throughout the entirety of the cell. The local electric- (CaT) and caffeine-evoked Ca^2+^ transient (cCaT) peak near the periphery (ss, 3 CRUs on each side) and central area (cc, 3 CRUs) were measured during the last 20 electric-evoked Ca^2+^ transients or caffeine stimulation and the ratio of central vs. surface electric-evoked Ca^2+^ transient peaks (cc/ss (ratio)) calculated from the average. The ratio of central vs. surface CaT and cCaT [Ca^2+^]_Cyto_ peaks (cc/ss (ratio)) are similar between experiment and simulation. **C)** Post-pacing caffeine-induced Ca^2+^ transient (cCaT) traces under 0.5 Hz voltage-clamp control in experimental (Voigt Niels et al., 2012) (i) and simulated (ii) cells. Voltage-clamp protocol is shown inset. Diastolic (iii) and systolic (iv) CaT [Ca^2+^], cCaT amplitude (v) and decay constant (vi) in experimental (Voigt Niels et al., 2012, 2014) and simulated cells with varying t-tubule densities (Dense, Median, Detubulated) as marked by respective blue, black, and orange bars.

To validate the role of TATS in subcellular Ca^2+^ signaling, we compared the spatial and temporal properties of Ca^2+^ sparks and waves in myocytes with and without tubules. Cells devoid of TATS exhibit a hallmark U-shaped Ca^2+^ wave, whereby Ca^2+^ rises at the cell periphery upon electric field stimulation and then propagates from the cell periphery to the center, as also shown in experimental linescan images (**Fig. 6Ai**, top) (Kirk *et al.*, 2003) and in our model (**Fig. 6Aii**, top). However, cells with central TATS exhibit W-shape Ca^2+^ waves (**Fig. 6Ai**, bottom) (Kirk *et al.*, 2003), indicating increased synchrony of Ca^2+^ release in inner areas where tubules are present. The simulation linescan images recapitulate the results of the experiment (**Fig. 6Aii**, bottom).

**Figure 6.**
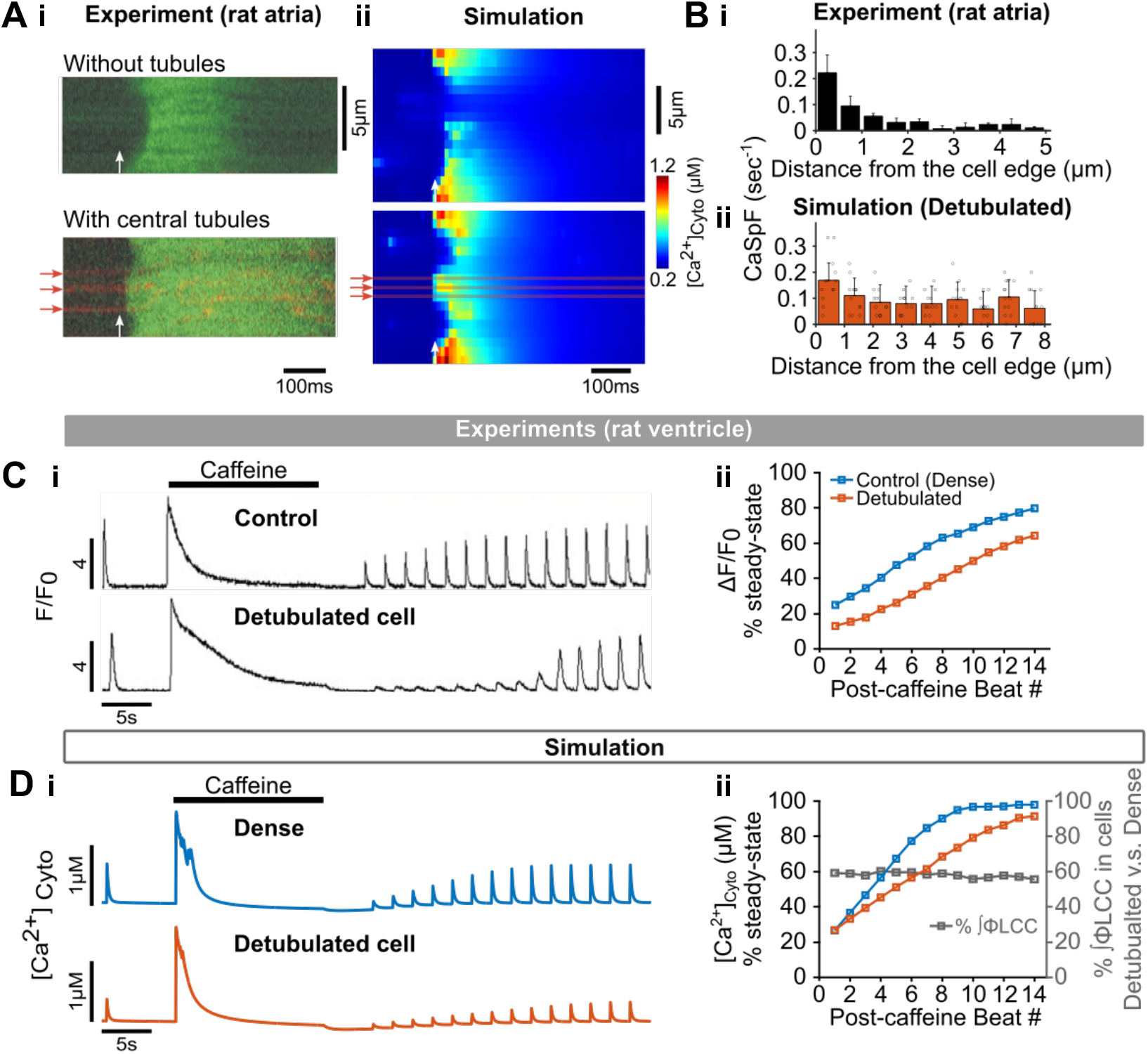
Validation of the effects of tubule-loss on Ca^2+^ signaling. **A)** Linescan images of experimental (Kirk et al., 2003) (i) and simulated (ii) atrial cells without (top) or with (bottom) tubules. Tubule locations are marked by red arrows and the stimulus timing is indicated with white arrows. The axial tubules in simulated cells are 3*3 CRUs in the center x-direction with whole-cell length. **B)** Spatial distribution of Ca^2+^ sparks in experimental rat atrial cells (Brette et al., 2005) (i) and simulated detubulated cells (ii). To replicate the experimental protocol, the simulated cell was paced at 0.5 Hz to steady state with a subsequent 30 s quiescence where Ca^2+^ sparks along 13 transversal scan lines (i.e., 13 y-axis lines equally distributed between 10^th^ and 46^th^ CRUs on the central x-axis) were measured and averaged. **C)** Representative CaT traces showing SR Ca^2+^ release following caffeine application and subsequent transient recovery (i) in experimental rat ventricular cells with dense tubules (top) and without tubules following detubulation (bottom) (Brette et al., 2005). Post-caffeine CaT recovery from experimental observation shows slower post-caffeine CaT recovery in detubulated cells (ii). **D)** Representative CaT traces showing SR Ca release following pseudo caffeine application (clamping RyR P_O_ at 0.2) and subsequent transient recovery (i) in simulated cells with dense tubules (top) and without tubules following detubulation (bottom). Simulated post-caffeine CaT recovery data (ii) is similar to experimental observation, in which the ratio of Ca^2+^ influx via LCC in detubulated vs. densely tubulated cells is reduced resulting in slower post-caffeine CaT recovery.

**Figure 7.**
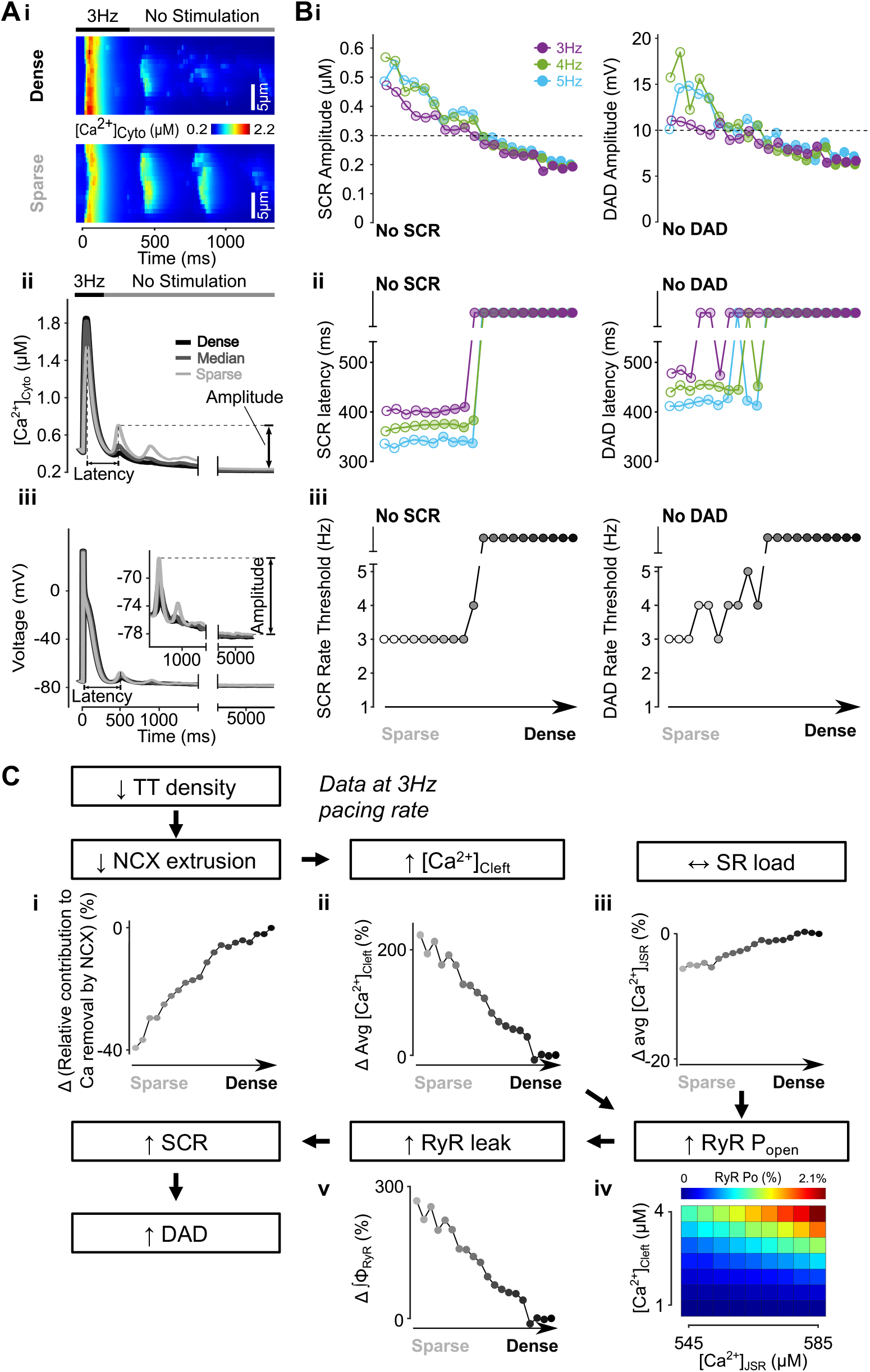
Loss of TATS promotes spontaneous Ca^2+^ release events (SCRs) and delayed after-depolarizations (DADs). **A)** Transverse line scan of cytosolic Ca^2+^ for cells with dense or sparse tubules (i), global cytosolic Ca^2+^ (ii) and voltage (iii) traces showing the final 3Hz-paced beat and subsequent no-stimulation period for observation of SCRs and DADs. SCR and DAD amplitude was calculated as the difference between the first peak maximum and the diastolic minimum in the no-stimulation period and latency as the duration between the peak of the last stimulated beat and the first peak of no-stimulation period. **B)** Amplitude (i), latency (ii) and rate threshold (iii) for SCRs and DADs with respect to increasing tubular density from detubulated to densely tubulated cells. Amplitudes over 0.3 μM and 10 mV were set as thresholds for SCRs and DADs respectively, with events under these cut-offs deemed as ‘no SCR’ or ‘no DAD’ and indicated by the dashed line. **C)** Mechanism underlying promotion of SCRs and DADs in cells with sparse tubules. Biomarkers were determined from the first 100 ms of the no-stimulation period and normalized to those of cells with dense tubules, with the Ca^2+^-dependence of RyR P_O_ determined by an in-silico bilayer study. In cells with fewer tubules, Ca^2+^ removal by NCX is reduced (i) leading to elevated cleft Ca^2+^ concentration (ii) but no change in SR load (iii). Increased cleft Ca^2+^ results in augmented RyR P_O_ (iv) and RyR leak (v) leading to increased SCRs and DADs.

**Figure 8.**
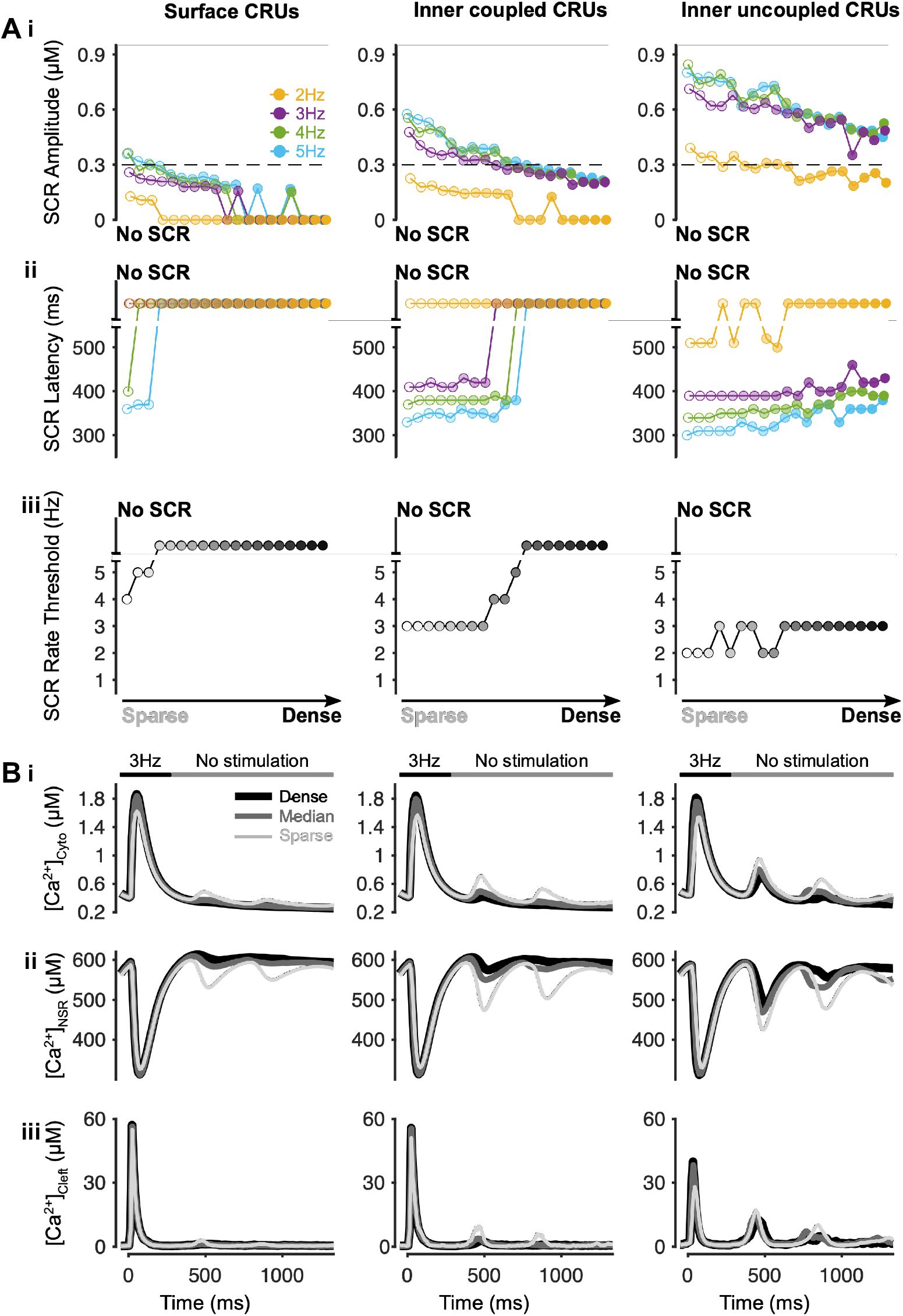
Promotion of SCRs is greatest in inner uncoupled CRUs in cells with fewer tubules. **A)** Amplitude (i), latency (ii) and rate threshold (iii) for local SCRs of surface (left), inner coupled (middle), and inner uncoupled (right) CRUs with respect to increasing tubular density. **B)** Ca^2+^ concentration in the cytosol (i), network SR (ii) and cleft (iii) of surface (left), inner coupled (middle) and inner uncoupled (right) CRUs following pacing at 3 Hz to examine SCRs.

To investigate the subcellular regional differences of Ca^2+^ sparks properties, Brette et. al. (Brette *et al.*, 2005) measured Ca^2+^ spark frequency (CaSpF) at 1 μm intervals from the cell edge, and found that diastolic Ca^2+^ sparks in rat atrial myocytes that lack TATS occurred predominantly at the cell periphery (**Fig. 6Bi**). Similarly, the CaSpF in the detubulated human atrial myocyte model was higher near the cell edge compared with the cell interior (**Fig. 6Bii**). In the same study, caffeine was applied to rat ventricular myocytes to empty the SR and monitor the gradual recovery of the Ca^2+^ transient. In these experiments the Ca^2+^ transient amplitude recovered more slowly in detubulated ventricular cells vs. control (**Fig. 6Ci**) (Brette *et al.*, 2005). The experimental summary data indicates that the number of beats required for 50% recovery was ~5.5 in control cells and ~9.9 in detubulated cells (**Fig. 6Cii**). The model recapitulates this behavior (**Fig. 6Di-ii**), in which loss of TATS slows post-caffeine Ca^2+^ transient amplitude recovery. Brette et. al. (Brette *et al.*, 2005) hypothesized that the slower recovery in detubulated cells is due to the reduced Ca^2+^ influx associated with TATS loss leading to slower SR refilling. In alignment with this hypothesis, our model shows that the Ca^2+^ influx via LCCs during the post-caffeine pacing period is reduced by 40% in the detubulated cell compared with the cell with dense TATS (**Fig. 6Dii** grey line). Therefore, simulated effects of human atrial myocyte detubulation on subcellular and whole-cell Ca^2+^ signaling mirror experimental observations in rat atrial and ventricular myocytes. Namely, our model recapitulates 1) the spatiotemporal characteristics of Ca^2+^ waves in the presence and absence of TATS, 2) the Ca^2+^ spark regional distribution in atrial myocytes, and 3) the time course of global [Ca^2+^]_Cyto_ transient amplitude recovery in the cells without and with TATS.

### Loss of TATS reduces NCX-mediated Ca^2+^ extrusion, elevating cleft Ca^2+^ and RyR P_O_ resulting in enhanced SCR events and promotion of DADs

With our validated model, we sought to investigate how changing TATS affect arrhythmic biomarkers, namely SCRs and DADs. We subjected our population of models with varying tubular structures and densities to pacing at various cycle lengths followed by a pause to detect any diastolic (unstimulated) activity. Representative linescan images from densely and sparsely tubulated myocyte models in **Fig. 7Ai** show that in the sparsely tubulated cell the AP-triggered Ca^2+^ wave was less synchronous, with larger and more frequent SCRs observed in the unstimulated period compared to the cell with dense TATS. Furthermore, the cell with sparse TATS displayed a smaller AP-triggered Ca^2+^ transient, but larger SCRs (**Fig. 7Aii**) and DADs (**Fig. 7Aiii**). These results were confirmed when repeating the stochastic simulations with different random seeds (**Fig. 9A**). Analysis of the whole myocyte model population with varying TATS revealed that cells with sparser TATS have larger SCRs and DADs (**Fig. 7Bi**), and a shorter latency of SCRs and DADs (**Fig. 7Bii**). Since SCRs and DADs are generally more likely to occur at increasing pacing rates, we measured the pacing rate threshold for SCRs and DADs, i.e., the slowest pacing rate at which these events occurred. Our model predictions indicate that the rate thresholds for SCRs and DADs are lower in cells with sparse vs. dense TATS (**Fig. 7Biii**), suggesting that cells with a low density of tubules may be more susceptible to Ca^2+^-driven arrhythmia. We further utilized the model to reveal the mechanisms by which loss of tubules promotes SCRs and DADs. We found that in cells with sparse TATS, Ca^2+^ extrusion by NCX is reduced (**Fig. 7Ci**) leading to elevated diastolic cleft Ca^2+^ concentration (**Fig. 7Cii**). The increased cleft Ca^2+^ results in augmented diastolic RyR P_O_ (**Fig. 7Civ**) and RyR leak (**Fig. 7Cv**) leading to increased SCRs and DADs. Conversely, the lack of appreciable changes in SR load (**Fig. 7Ciii**) did not strongly affect RyR P_O_. Overall, our simulations suggest that loss of TATS is associated with reduced NCX-mediated Ca^2+^ extrusion, which alters RyR function and promotes SCRs and DADs.

**Figure 9.**
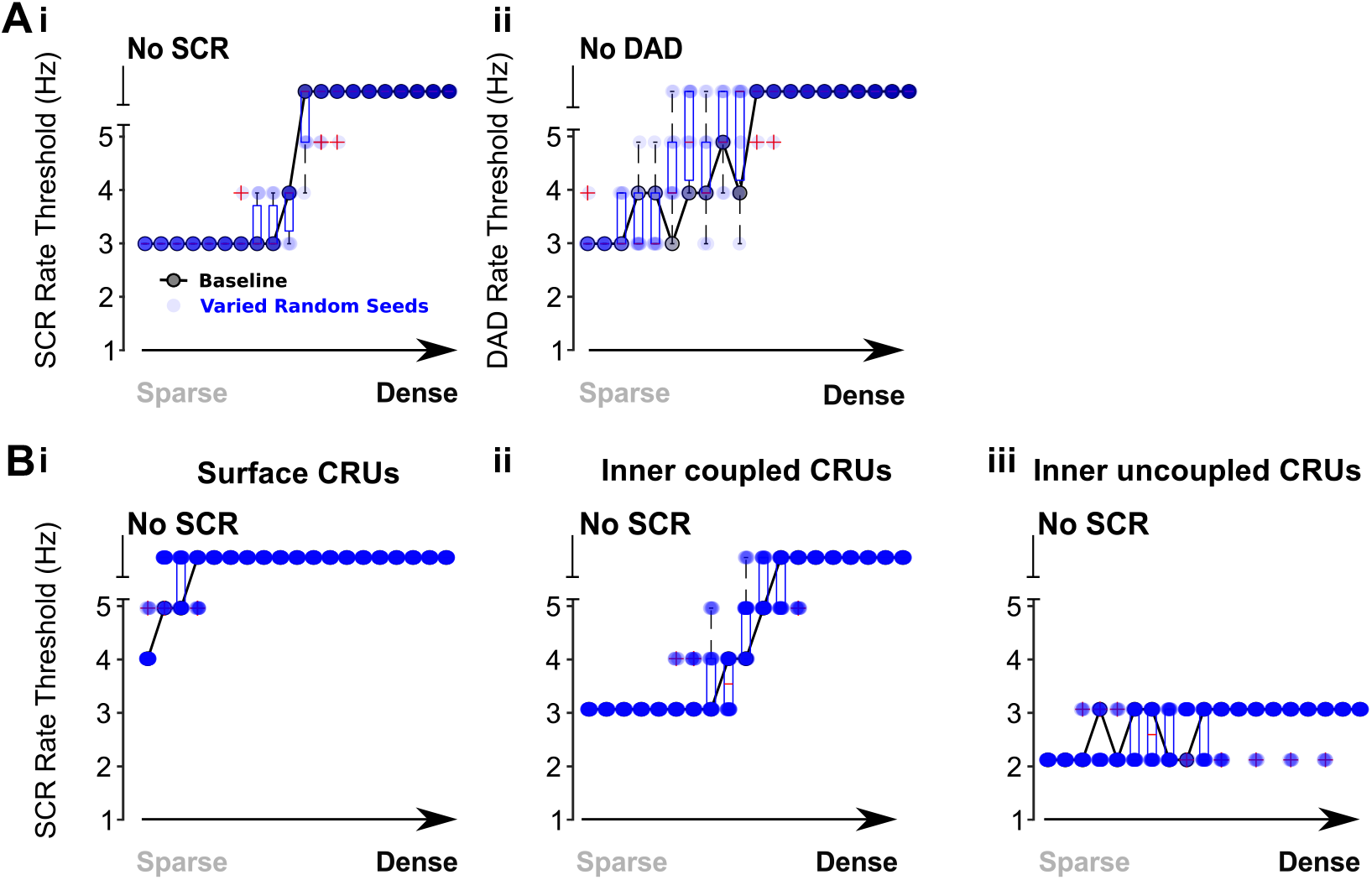
Varying random seeds regulating stochasticity does not change the effects of TATS loss on SCRs and DADs. **A)** Rate threshold for SCRs (i) and DADs (ii) with respect to increasing tubular density from detubulated to densely tubulated, with 10 different random seeds (blue). The baseline results are superimposed. **B)** TATS-loss-induced promotion of SCRs is greatest in inner uncoupled CRUs in cells with fewer tubules in all three conditions of varying uncoupled CRU compartments.

### SCR properties vary between spatially distinct CRUs, with SCR threshold and latency decreased and amplitude increased in inner uncoupled CRUs

We further sought to determine whether regional variations exist in the effects of varying tubular density on the latency, amplitude, and pacing threshold for SCRs. To do so, we analyzed the averaged Ca^2+^ transient of surface CRUs, inner CRUs coupled to TATS, and inner CRU that are not coupled to TATS in each cell in the population with varying TATS and quantified the biomarkers of the local Ca^2+^ transients. We found that the amplitude of SCRs is larger (**Fig. 8Ai**) and the latency reduced (**Fig. 8Aii**) in inner uncoupled CRUs, where the pacing threshold for SCR is lower (**Fig. 8Aiii** and **9B**) compared to inner coupled and surface CRUs. The representative traces of average cytosolic, network SR, and cleft Ca^2+^ concentration following pacing at 3 Hz (**Fig. 8B**) also show larger SCRs in inner uncoupled CRUs independent of the TATS density. The simulation results indicate that while loss of TATS promotes SCRs throughout the whole cell, uncoupled CRUs may play a more important role in SCR incidence vs. inner coupled CRUs and surface CRUs.

## Discussion

In this study we built, parameterized, and validated a three-dimensional model of the human atrial myocyte, coupling electrophysiology and Ca^2+^ handling with subcellular spatial details governed by the TATS. This work quantitatively explains how TATS loss disrupts diastolic Ca^2+^ homeostasis and electrophysiological stability in human atrial cells. Specifically, we demonstrated that TATS loss reduces NCX-mediated Ca^2+^ removal leading to increased cleft Ca^2+^ concentration that promotes SCRs (**Fig. 7C**), especially in inner CRUs (**Fig. 8**), and subsequent DADs (**Fig. 7C**). These findings provide mechanistic insight into how atrial TATS remodeling can lead to Ca^2+^-driven proarrhythmic behavior in the physiological range that may ultimately contribute to the arrhythmogenic state in both HF and AF.

### TATS loss is associated with altered Ca^2+^ homeostasis

Previous experimental studies have shown atrial TATS loss is associated with disrupted Ca^2+^ handling in paroxysmal AF (Wakili *et al.*, 2010), persistent AF (Lenaerts *et al.*, 2009), and HF induced by both rapid ventricular pacing (Dibb *et al.*, 2009) and myocardial infarction (Kettlewell *et al.*, 2013), as also shown in the ventricle in HF (Balijepalli *et al.*, 2003; Louch *et al.*, 2004, 2006; Cannell *et al.*, 2006). The remodeling of the TATS in disease has two main functional consequences: (i) cell membrane channel localization and currents are altered and (ii) RyRs are uncoupled from LCCs and other transporters in the sarcolemma. In both atrial and ventricular myocytes, loss of TATS causes decreased LCC Ca^2+^ influx, accompanied by shortening of APD and thus refractory period that can increase vulnerability to arrhythmic activity (Bosch *et al.*, 1999; Kneller *et al.*, 2002; Brette *et al.*, 2006; Lenaerts *et al.*, 2009; Wakili *et al.*, 2010). These findings were reproduced in our simulation results, where detubulated cells had shorter APD and reduced Ca^2+^ influx compared to those with denser TATS (**Fig. 4Cii** and **6Dii**). In addition to I_Ca_ and APD changes, TATS loss increases the number of uncoupled RyRs (Song *et al.*, 2006) resulting in asynchronous CICR during the systolic period (Song *et al.*, 2006; Louch *et al.*, 2006), and damaged excitability and contractility (Sacconi *et al.*, 2012). Together, the compounding effects of reduced Ca^2+^ influx and RyR coupling present in the diastolic period, with slowed post-caffeine Ca^2+^ SR reloading, reduced diastolic Ca^2+^ spark frequency (Brette *et al.*, 2005), and enhanced SCRs in uncoupled CRUs in HF (Dries *et al.*, 2018), as recapitulated in our model.

Similar to TATS loss in disease, asynchrony of CICR is also reported in various experiments of atrial cells with sparse TATS where many central RyRs are uncoupled. Here, while triggered release occurs around the cell periphery, central CICR occurs as a U-shaped wave of propagation leading to delayed and impaired Ca^2+^ release in the cell interior (Kirk *et al.*, 2003; Woo *et al.*, 2005; Dibb *et al.*, 2009; Lenaerts *et al.*, 2009; Smyrnias *et al.*, 2010; Wakili *et al.*, 2010; Frisk *et al.*, 2014; Yue *et al.*, 2017). The asynchronous CICR in atrial myocytes leads to smaller Ca^2+^ transient amplitude and longer rising duration, which is linked with weaker contractility compared to ventricular myocytes (Tanaami *et al.*, 2005; Narolska *et al.*, 2005; Walden *et al.*, 2009; Smyrnias *et al.*, 2010). Interestingly in atrial cells with extensive TATS, TAT-coupled RyRs are hyperphosphorylated which facilitates enhanced Ca^2+^ release and promotes the propagation of Ca^2+^ waves (Brandenburg *et al.*, 2016). As such, loss of TATS in these cells would disrupt both triggered and propagated Ca^2+^ release diminishing systolic contractility. In addition to the effect on Ca^2+^ transient amplitude, computational studies have predicted that TATS loss also enhances alternans susceptibility (Li *et al.*, 2012; Nivala *et al.*, 2015) and the likelihood of triggered Ca^2+^ waves during the AP due to the latency of Ca^2+^ release by the large pool of orphaned RyRs following LCC opening (Shiferaw *et al.*, 2017, 2018, 2020).

### Reduced local NCX activity underlies diastolic Ca^2+^ and V_m_ instabilities associated with TATS loss

While systolic Ca^2+^ abnormalities due to TATS loss have been extensively characterized, the impact of TATS remodeling on Ca^2+^ homeostasis during the diastolic period is not fully understood. Though much consideration has been given to the impact of altered LCC localization, changes in other tubular proteins, namely NCX, likely play a key role. Indeed, we predict that the effect of TATS loss on NCX function underlies diastolic Ca^2+^ and voltage instabilities. Experimental observations indicate TATS loss disrupts NCX function to promote SCRs in atrial myocytes, with preliminary data from computational studies predicting the association between the few TATS and SCRs (Colman *et al.*, 2016). Both NCX and PMCA are highly expressed within the TATS (Despa *et al.*, 2003; Chase & Orchard, 2011; Schulson *et al.*, 2011; Swift *et al.*, 2012) and NCX on TATS has been shown to co-localize with RyRs (Thomas *et al.*, 2003; Jayasinghe *et al.*, 2009; Schulson *et al.*, 2011; Biesmans *et al.*, 2011). Loss of TATS in NCX knock-out has also been shown to be associated with abnormal Ca^2+^ cycling that impacts contractility and rhythm (Yue *et al.*, 2017). As such, TATS loss induces 1) decreased NCX expression and subsequent reduced local Ca^2+^ extrusion, 2) NCX-RyR decoupling, and localized inner subcellular Ca^2+^ accumulation. Our results support this mechanism and demonstrate that disrupted NCX-mediated Ca^2+^ removal caused by TATS loss contributes to SCRs by elevating [Ca^2+^]_Cleft_ and subsequently enhancing RyR P_O_ (**Fig. 7C**), especially in inner uncoupled CRUs (**Fig. 8A**). From this point of view, though the NCX upregulation that occurs in AF eventually enhances Ca^2+^-voltage instability (Voigt *et al.*, 2012), it may act to limit enhanced SCRs and shortened APD responses to TATS loss and reduced I_Ca_ and therefore be an initial compensatory response. Increased TATS density with enhanced NCX expression in SERCA-KO mice is also believed to be a compensatory response (Swift *et al.*, 2012). While under these conditions reduced NCX contributes to a pro-arrhythmic state, the opposite can also be true whereby altered NCX function can shift the balance between SCRs and Ca^2+^-voltage instability coupling in an anti-arrhythmic manner (Antoons *et al.*, 2012). Though it is difficult to isolate the independent role of decreased NCX from reduced I_Ca_ and orphaning of RyRs that also occur with TATS loss, Ca^2+^ signal silencing and reduced central Ca^2+^ transient that may protect the cell from SCRs have been reported experimentally in atrial myocytes following 5-7 days of rapid atrial pacing (Wakili *et al.*, 2010; Greiser *et al.*, 2014). It is unclear whether this short-term protective effect of TATS loss is transient and whether the longer-term remodeling that occurs, for example in the transition between paroxysmal to chronic AF, leads to a swing towards the pro-arrhythmic state (Lenaerts *et al.*, 2009; Wakili *et al.*, 2010). Given our simulations of TATS loss and the resultant reduced NCX-mediated Ca^2+^ extrusion cause increased SCRs and DADs, we suggest that ultimately the pro-arrhythmic increase in [Ca^2+^]_Cleft_ overrides any benefit of reduced triggered central Ca^2+^ release. This tool can be used to investigate transitional remodeling that occurs during disease progression.

### Arrhythmogenic waves in human atrial myocytes originate from the inner uncoupled CRUs

In addition to investigating the impact of variable TATS densities on the occurrence of SCRs and DADs, through incorporating different spatial CRU domains our model has permitted examination of the origin of these events. Experimental studies have previously shown diastolic Ca^2+^ sparks occur primarily around the cell periphery in atrial cells with no TATS (Brette *et al.*, 2005), suggesting that SCRs initiate in coupled CRUs. Interestingly, we observed this in the model at slow pacing rates (**Fig. 6B**), but with rapid pacing, as seen in atrial tachycardia and AF, SCRs were larger and showed greater incidence in inner uncoupled CRUs vs. coupled CRUs (**Fig. 8**). We suggest that these regional differences are indeed dependent on the stimulation rate and, notably, they are similar to those observed in HF ventricular myocytes, where spark frequency increases in uncoupled vs. coupled release sites with increasing pacing frequency (Dries *et al.*, 2018).

At faster rates, the increase in SR Ca^2+^ loading and thus the increase in RyR leak (Shiferaw *et al.*, 2017) impact more strongly the inner uncoupled CRUs by enhancing SCRs, while local NCX Ca^2+^ extrusion eases Ca^2+^ sparks in coupled CRUs. Compared to coupled CRUs where Ca^2+^ is rapidly extruded by colocalized NCX thus limiting the magnitude of SCRs, Ca^2+^ leak from uncoupled CRUs is not readily removed from the cytosol and must diffuse to be extruded by NCX. As such, the inner Ca^2+^ concentration is relatively higher than that at the periphery. In addition to changes in SR load, fast pacing elevates inner diastolic [Ca^2+^]_Cleft_ causing increased RyR P_O_ and thus greater leak in inner uncoupled CRUs vs. surface CRUs due to spatial differences in NCX extrusion (**Fig. 7C**). This is in contrast to that at slower pacing rates where lower diastolic [Ca^2+^]_Cleft_ results in similar RyR P_O_ in coupled vs. uncoupled CRUs (**Fig. 2Bii** and **7Civ**). The higher simulated RyR P_O_ near the cell periphery vs. interior at lower pacing rates (**Fig. 6B**) is induced by the differences in Ca^2+^ diffusion characteristics at surface vs. inner CRUs and the presence of background Ca^2+^ currents on the external sarcolemma of detubulated cells.

In addition, varying TATS density does not appreciably change the SCR rate threshold in surface and inner uncoupled CRUs, with surface CRUs being generally more stable and inner uncoupled CRUs typically being more prone to SCR. Instead, varying TATS changes the balance between (stable) coupled and (unstable) uncoupled CRUs to determine the net effect for SCR, with the most remarkable changes seen in the TATS-dependence of SCR in inner coupled CRUs. Given that the extensive loss of TATS in HF and AF consequently increases the number of inner uncoupled CRUs, this may contribute to the increased arrhythmic activity observed in these diseases.

### Model assumptions and limitations

We present, for the first time, a fully integrated human atrial ECC model that is coupled with an experimentally informed population of TATS structures. We incorporated TATS properties measured in an experimental manner to ensure the TATS population replicates experimental observations (**Fig. 1Bi-ii**) and allows for virtual-patient population and high-throughput simulation in future TATS studies. Model parameterization (**Fig. 1, 2A-B, 3**) and validation (**Fig. 2C, 4-6**) utilized extensive human-specific independent datasets to generate human-specific results and conclusions that are highly translatable. As such, using this model we have systematically evaluated the role of TATS disruption in Ca^2+^-driven proarrhythmic behavior in the physiological range.

We acknowledge that other factors may contribute to the effect of varying TATS on local and global Ca^2+^ signaling, including variation in the volume of the release sites, non-uniform distribution of Ca^2+^ handling proteins (Herraiz-Martínez *et al.*, 2022), and alterations in their regulatory state (Brandenburg *et al.*, 2016), as discussed below. In our compartmental model, RyRs are sensitive to the rapid changes in cleft Ca^2+^ favored by the narrow cleft (i.e., the surrounding area near RyRs), which is critical for both CICR and Ca^2+^ sparks. While our model assumes that all coupled and uncoupled CRUs have the same Ca^2+^compartments, as also done in previous computational studies (Shiferaw *et al.*, 2017, 2020; Song *et al.*, 2018), it is conceivable that RyRs release Ca^2+^ in a larger compartment in CRUs that are not coupled with T-tubules. To address this possibility, we assessed the impact of altered Ca^2+^ compartmentation in orphaned/uncoupled CRUs. Namely, we performed simulations in which uncoupled CRUs had increased (+50%) submembrane volume (v_s_), increased cleft volume (v_Cleft_), or faster Ca^2+^ diffusion between cleft and submembrane (*τ*_Cleft↔s_ decreased by 50%). In all these simulated conditions, our main conclusions on the local and global effects of TATS loss on SCR (and DAD) remained unaffected (**Fig. 10**). Additionally, we note that Ca^2+^ diffusion out of the cleft (*τ*_Cleft↔s_ ~0.022 ms, estimated by (Restrepo *et al.*, 2008)) is much faster than RyR opening and closure (0.7~1.9 ms) (Györke & Györke, 1998). Thus, sustained RyR flux due to longer channel openings (or fewer closures), dictated by RyR Ca^2+^ sensitivity and unitary flux, is most potent for maintaining high local Ca^2+^.

**Figure 10.**
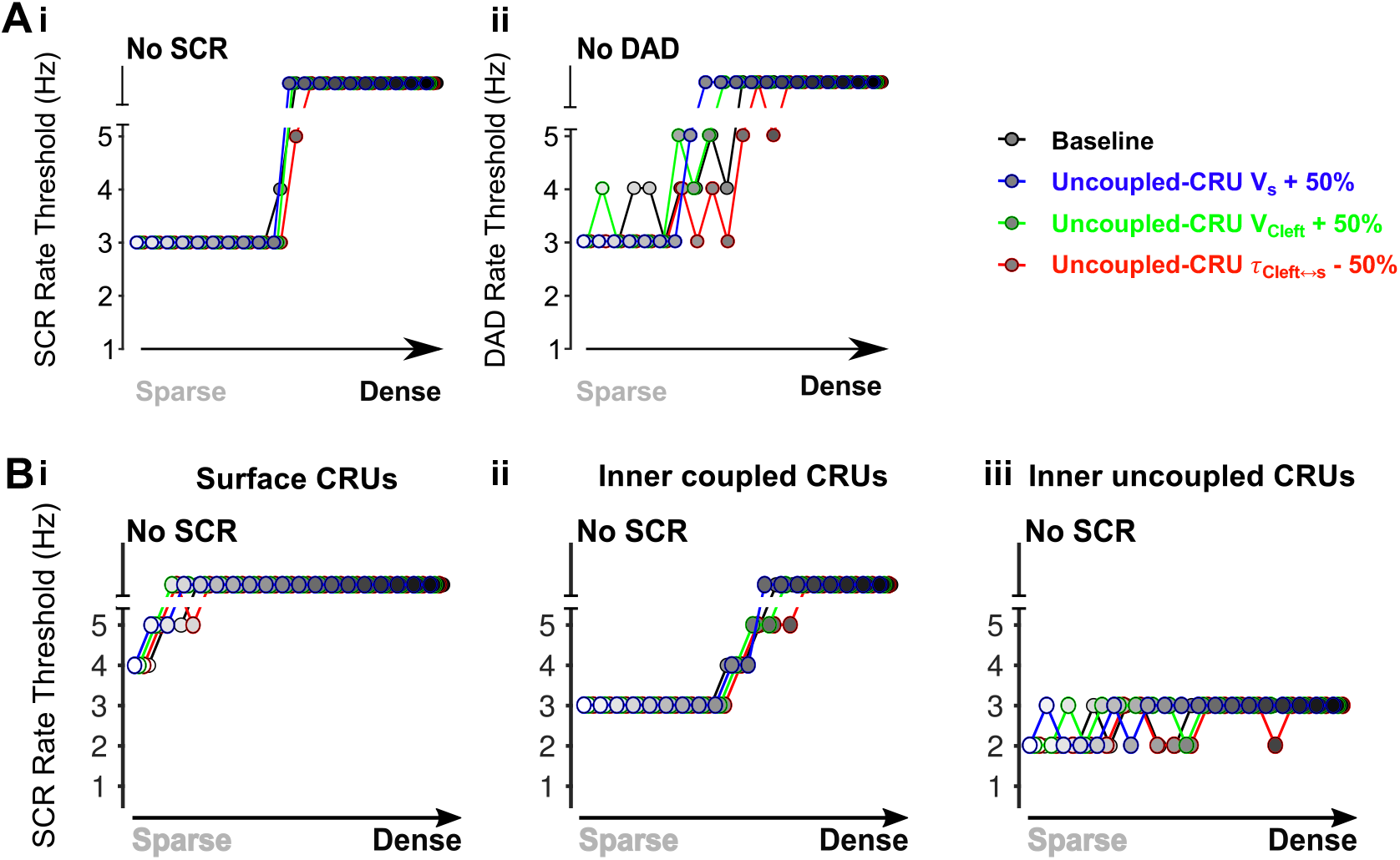
Varying compartmentalization of uncoupled CRUs does not change the effects of TATS loss on SCRs and DADs. **A)** Rate threshold for SCRs (i) and DADs (ii) with respect to increasing tubular density from detubulated to densely tubulated. Uncoupled CRU compartments retained the maximum ion flux strength but had 1) submembrane volume (v_s_) increased by 50% (blue), 2) cleft volume (v_Cleft_) increased by 50% (green), or 3) the time scale of Ca^2+^ diffusion between cleft and submembrane (*τ*_Cleft↔s_) decreased by 50% (red). **B)** TATS-loss-induced promotion of SCRs is greatest in inner uncoupled CRUs in cells with fewer tubules in all three conditions of varying uncoupled CRU compartments.

Our model assumes that all CRUs are composed of the same ensemble of ion channels and transporters, with the only differences being incorporated between CRUs that are coupled vs. uncoupled to the cell membrane. Nevertheless, heterogeneity has been reported in the distribution of Ca^2+^-handling proteins (e.g., CSQ and RyR) in human atrial myocytes (Herraiz-Martínez *et al.*, 2022). The effect of altering the expression and distribution of key Ca^2+^ handling proteins (i.e., NCX, RyR, and CSQ) on SCR and arrhythmogenic outcomes is the focus of a companion paper (Zhang et al).

Similarly, our model does not explicitly incorporate the mechanisms by which PKA and CaMKII influence the function of LCC, RyR, and PLB in normal physiology nor their regulatory state in the face of disease-induced ultrastructural remodeling. For example, RyR clusters at axial tubule-SR junctions are hyperphosphorylated to facilitate spontaneous macro Ca^2+^ sparks (duration and width) and rapid AP-evoked Ca^2+^ signals in mouse atria (Brandenburg *et al.*, 2016). A computational study predicted that this local hyperphosphorylation may increase the incidence at these sites but reduce the size of the SCRs (Sutanto *et al.*, 2018). However, the latency difference of AP-evoked Ca^2+^ release between axial tubule coupled junctions and those at the surface is much more pronounced in mouse and rat atrial myocytes but less evident in the more human-like rabbit atrial myocyte (Brandenburg *et al.*, 2018). Such difference suggests the effects of RyR hyperphosphorylation differs between species. We carried out additional simulations in which hyperphosphorylation of the TATS-associated RyRs was modeled by increasing the Ca^2+^ sensitivity of RyR P_O_ (i.e., reducing kcp) (**Fig. 2Bii-iii**). Our results indicated that this differential regulation at the TATS-associated RyRs is not necessary to explain the faster AP-evoked Ca^2+^ release at the axial tubule coupled junctions vs. surface region (**Fig. 11Aii-iii**) shown in rat atrial myocyte experiments (**Fig. 11Ai**) (Brandenburg *et al.*, 2016). In these experiments, higher LCC density in axial tubule junctions is reported (Brandenburg *et al.*, 2016), which may further contribute to the regional differences in AP-evoked Ca^2+^ signaling in rat atrial myocytes. On the other hand, our simulations also indicate that the presence of axial tubules can in itself lead to higher Ca^2+^ spark frequency near axial tubules (**Fig. 11Bii**), with hyperphosphorylation of TATS-associated RyRs only modestly further increasing the Ca^2+^ spark frequency at these sites (**Fig. 11Biii**), similar to experimental results (**Fig. 11Bi**) (Brandenburg *et al.*, 2016). When simulating the effects of varying TATS density in the presence of hyperphosphorylation at the TATS-associated RyRs, we observed that while TATS loss still promotes SCRs and DADs (**Fig. 11CD**), cells with denser TATS exhibited a lower rate threshold for SCRs and DADs compared with our baseline model (**Figs. 7 and 11C**). On the other hand, in ventricular myocytes with HF-induced TATS loss, non-coupled RyRs are more sensitive to CaMKII-inhibition, which suggests CaMKII may preferentially phosphorylate RyRs in uncoupled vs. coupled CRUs to initiate SCRs in the uncoupled CRUs (Dries *et al.*, 2018). As such, CaMKII hyperphosphorylation of uncoupled RyRs may ensue to compensate for lower systolic Ca^2+^ release in uncoupled CRUs in atrial myocytes, with the maladaptive effect to increase diastolic RyR leak. Simulating hyperphosphorylated RyRs in uncoupled CRUs does indeed predict enhanced diastolic [Ca^2+^]_Cyto_, especially in uncoupled CRUs (not shown).

**Figure 11.**
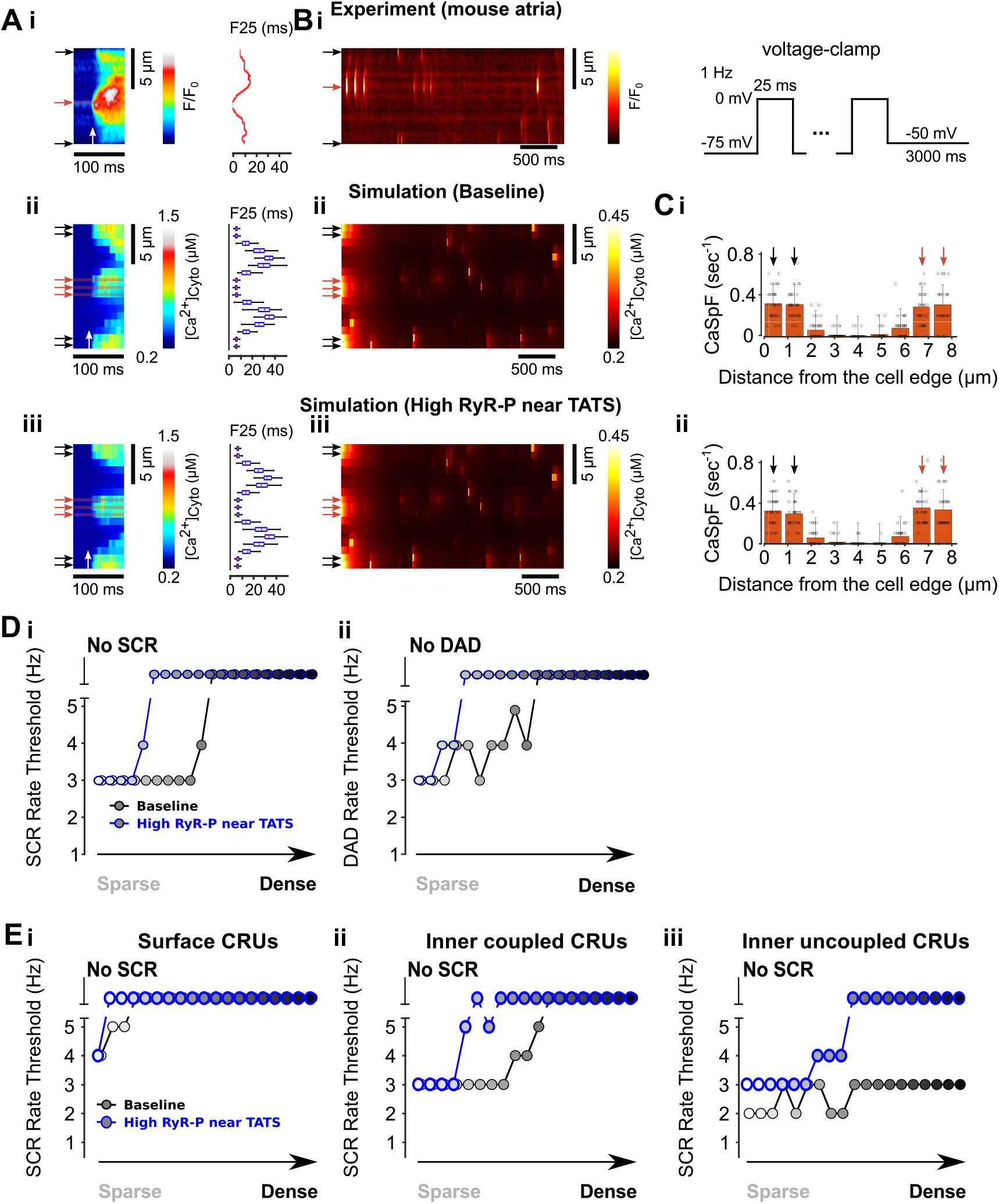
TATS-associated RyR hyperphosphorylation does not change the effects of TATS loss on SCRs and DADs. **A)** Experimental observations in a mouse atrial myocyte (i) showing the Ca^2+^ transient near TATS junctions (red arrow) vs. surface (black arrow) vs. uncoupled areas following stimulation (white arrow). The time to reach 25% of subcellular Ca^2+^ transient amplitude (F25) is reduced at TAT junctions suggesting Ca^2+^ release at TATS junctions occurs more rapidly following electrical stimulation (Brandenburg et al., 2016). Simulation of the model with axial tubules replicated the rapid response of subcellular Ca^2+^ signaling near tubules and surface without significant difference in F25 between surface and tubules (ii). This lack of difference remained unaltered with hyper-phosphorylation of TATS-associated RyRs (iii). **B)** During the voltage-holding period following 1-Hz voltage-clamp (protocol as an insert), Ca^2+^ spark occurrence is increased near tubules (red arrows) vs. other areas experimentally in mouse atrial myocytes (Brandenburg et al., 2016) (i) and in the baseline (ii) and hyperphosphorylated TATS-associated RyR (iii) model simulation. **C)** Quantitative measurement of regional differences in Ca^2+^ spark frequency confirming increased spark occurrence at TATs in the baseline (i) and hyperphosphorylated RyR (ii) simulations. **D)** Relationship between SCR (i) and DAD (ii) occurrence in the baseline and hyperphosphorylated simulations with respect to increasing tubular density from detubulated to densely tubulated cells. Similar to the baseline model, TATS loss continues to reduce the rate threshold of SCRs (i) and DADs (ii) in the hyperphosphorylated RyR simulation, however this occurs at a lower TAT density. This is induced by lower SR load with leakier RyRs, which is in accordance with the conclusions in the companion paper (Zhang et al). **E)** SCR occurrence in surface (i), inner coupled (ii) and inner uncoupled (iii) CRUs with respect to increasing tubular density from detubulated to densely tubulated cells. When all RyRs in inner coupled CRUs are hyperphosphorylated, TATS-loss-induced promotion of SCRs in inner coupled and uncoupled CRUs remains apparent as in the baseline simulation. However, the threshold for SCRs is higher in following hyperphosphorylation with reduction in SCR threshold requiring more extensive TAT loss vs. baseline.

## Conclusions

We developed a computational platform to study the interaction between changes in TATS organization/density, Ca^2+^ handling, and electrophysiology in human atrial myocytes. Our model explains the mechanisms by which atrial myocytes with sparse TATS exhibit greater vulnerability to SCRs and DADs and provides insight into ionic and Ca^2+^ handling remodeling that occurs alongside TATS loss in disease.

## Abbreviations

AF: atrial fibrillation
AP: action potential
APD: AP duration
BCL: basic cycle length
CaT: cytosolic Ca^2+^ transient
CaTD_80_: CaT duration at 80% systolic level
CC: cell central area
cCaT: caffeine-induced CaT
CICR: Ca^2+^-induced Ca^2+^ release
CRU: Ca^2+^ release unit
CSQ: calsequestrin
Cyto: Cytosol
DAD: delayed afterdepolarization
d[Ca^2+^]_Cyto_/dt: derivative of CaT
dV/dt_max_: maximum upstroke velocity of depolarization
ECC: excitation-contraction coupling
HF: heart failure
i_Ca_: LCC unitary Ca^2+^ current
I_Ca_: LCC current
I_Cabk_: background Ca^2+^ current
i_Ca,K_: LCC unitary K^+^ current
i_Ca,Na_: LCC unitary Na^+^ current
I_ClCa_: Ca^2+^-activated Cl^-^ current
I_K1_: inward rectifier K^+^ current
I_K2P_: 2-pore-domain K^+^ current
I_K,Ach_: acetylcholine-activated K^+^ current
I_Kr_: rapidly activating delayed rectifier K^+^ current
I_KS_: slowly activating delayed rectifier K^+^ current
I_Kur_: ultrarapid delayed rectifier K^+^ current
I_leak_: SR leak current
I_Na_: fast Na^+^ current
I_Nabk_: background Na^+^ current
I_NaK_: Na^+^/K^+^ pump current
I_NCX_: Na^+^-Ca^2+^ exchanger current
I_PMCA_: PMCA current
I_rel_: RyR Ca^2+^ release
I_SK_: small-conductance Ca^2+^-activated K^+^ current
I_to_: transient outward K^+^ current
I_up_: SERCA uptake current
JSR: junctional SR
LCC: voltage-gated L-type Ca^2+^ channel
NCX: Na^+^-Ca^2+^ exchanger
NSR: network SR
PMCA: plasma membrane Ca^2+^ ATPase
P_O_: open probability
RMP: resting membrane potential
RyR: ryanodine receptor
SCR: spontaneous Ca^2+^ release
SERCA: SR Ca^2+^-ATPase
SR: sarcoplasmic reticulum
SS: cell surface area
TATS: transverse-axial tubular system
TT: Transverse tubule
V_m_: membrane voltage

